# Tonic Meningeal Interleukin-10 Upregulates Delta Opioid Receptor to Prevent Relapse to Pain

**DOI:** 10.1101/2023.06.08.544200

**Authors:** Kufreobong E. Inyang, Jaewon Sim, Kimberly B. Clark, Geron Matan, Karli Monahan, Christine Evans, Po Beng, Jiacheng “Vicky” Ma, Cobi J. Heijnen, Robert Dantzer, Gregory Scherrer, Annemieke Kavelaars, Matthew Bernard, Yasser Aldhamen, Joseph K. Folger, Geoffroy Laumet

## Abstract

Chronic pain often alternates between transient remission and relapse of severe pain. While most research on chronic pain has focused on mechanisms maintaining pain, there is a critical unmet need to understand what prevents pain from re-emerging in those who recover from acute pain. We found that interleukin (IL)-10, a pain resolving cytokine, is persistently produced by resident macrophages in the spinal meninges during remission from pain. IL-10 upregulated expression and analgesic activity of δ-opioid receptor (δOR) in the dorsal root ganglion. Genetic or pharmacological inhibition of IL-10 signaling or δOR triggered relapse to pain in both sexes. These data challenge the widespread assumption that remission of pain is simply a return to the naïve state before pain was induced. Instead, our findings strongly suggest a novel concept that: remission is a state of lasting pain vulnerability that results from a long-lasting neuroimmune interactions in the nociceptive system.

## Introduction

Chronic pain affects 20% of the population (Zelaya et al. 2020). Chronic pain does not necessarily develop as a continuation of acute pain but can develop asymptomatically long after apparent remission of the pain initiated by the primary insult. Additionally, 30-50% of patients with chronic pain exhibit relapsing pain (Hancock et al. 2015). Relapse to pain is observed in animal models and clearly involves mu-opioid receptors (µOR) (Campillo et al. 2011, Corder et al. 2013, Inyang et al. 2021). Indicating that ongoing suppression of a sensitized system is required to maintain remission from pain.

Communication between sensory neurons and immune cells contributes to the development of chronic pain (Shepherd et al. 2018, Ray et al. 2021, Szabo-Pardi et al. 2021) and recently we and others showed that such communication also contributes to the resolution of pain (Wei et al. 2019, Laumet et al. 2020, Parisien et al. 2022). A resolving role of immune cells is emerging in human as well (Parisien et al. 2022). Particularly, endogenous interleukin-10 (IL-10) is critical for the resolution of various chronic pain models (Krukowski et al. 2016, Song et al. 2016, Durante et al. 2021). In human, higher levels of IL-10 are reported in painless neuropathy compared to painful neuropathy (Uçeyler et al. 2007) and reduced IL-10 expression is observed in painful pancreatic and prostate cancer (Faupel-Badger et al. 2008, Wangzhou et al. 2021). Exogenous administration of IL-10 alleviates pain hypersensitivity in various preclinical models (Krukowski et al. 2016, Watkins et al. 2020, Prado et al. 2021). In addition to its inhibiting effects on proinflammatory immune cells (Donnelly et al. 1999, Moore et al. 2001), we recently showed that activation of IL-10 receptor, on sensory neurons contribute to the resolution of pain hypersensitivity induced by chemotherapy by normalizing neuronal excitability (Laumet et al. 2020) and similar findings were confirmed in preclinical migraine model (Guo et al. 2022). Given the critical role of IL-10 for the resolution of pain, we hypothesized that IL-10 is necessary for preventing relapse to pain.

Here, we found that IL-10 is produced by spinal meningeal immune cells and necessary to maintain remission from pain. RNA sequencing analysis revealed that IL-10 controls the expression of δ-opioid receptors (δOR) in the DRG and inhibition of δOR signaling triggered relapse to pain. Moreover, δOR analgesic activity increased during remission, but it is blocked by deletion of IL-10R1 on DRG neurons.

## Results

### IL-10 signaling prevents relapse to pain

Female and male mice treated with cisplatin (2 mg/kg x 3 consecutive days) developed mechanical hypersensitivity, that peaked at 4-8 days and reached remission by day 30 (**Figure 1A**). The resolution of cisplatin-induced mechanical hypersensitivity was impaired in absence of IL-10 (**Figure 1B**), consistent with the critical role of IL-10 for the resolution across pain models (Bang et al. 2006, Krukowski et al. 2016, Laumet et al. 2020, Wang et al. 2020, Singh et al. 2022). To test whether IL-10 is required to maintain remission from pain, on day 34, IL-10 signaling was blocked by intrathecal injection of neutralizing anti-IL-10 antibody. Inhibition of spinal IL-10 signaling reinstated pain hypersensitivity in cisplatin-treated female and male mice but had no effect on PBS-treated control mice (**Figure 1C, S1**). Additionally, inhibition of IL-10 signaling 3 months after cisplatin injection still reinstated pain hypersensitivity (**Figure 1D**). To determine if the reinstatement of pain behaviors by inhibition of IL-10 signaling is specific to chemotherapy-induced neuropathic pain, we used the incision postoperative pain model (Inyang et al. 2019, Inyang et al. 2021). Following remission of pain hypersensitivity, intrathecal injection of neutralizing anti-IL-10 antibody reinstated pain hypersensitivity in operated mice (**Figure 1E**). Because previous studies showed an essential role of IL-10R1 on DRG neurons for the remission of pain hypersensitivity (Laumet et al. 2020, Prado et al. 2021), HSV-Cre-mCherry viral vectors were injected into the hind paws of *Il10ra*^floxflox^ mice to downregulate IL-10R1 on DRG neurons. HSV virus have a tropism for sensory neurons (Hafezi et al. 2012) and infect neurons retrogradely via their axon terminals. Indeed, intraplantar injection of HSV viral vectors are known to reach DRG neurons (Wang et al. 2013, Willemen et al. 2018). The expression of mCherry was confirmed in the DRG but not in the spinal cord (**Figure S1B**). ∼50% of DRG neurons expressed mCherry after injection of HSV-Cre-mCherry viral vectors (**Figure S1C**) Downregulation of IL-10R1 on DRG neurons, reinstated mechanical pain hypersensitivity in cisplatin-treated mice in remission (**Figure 1F**).

**Figure 1.**
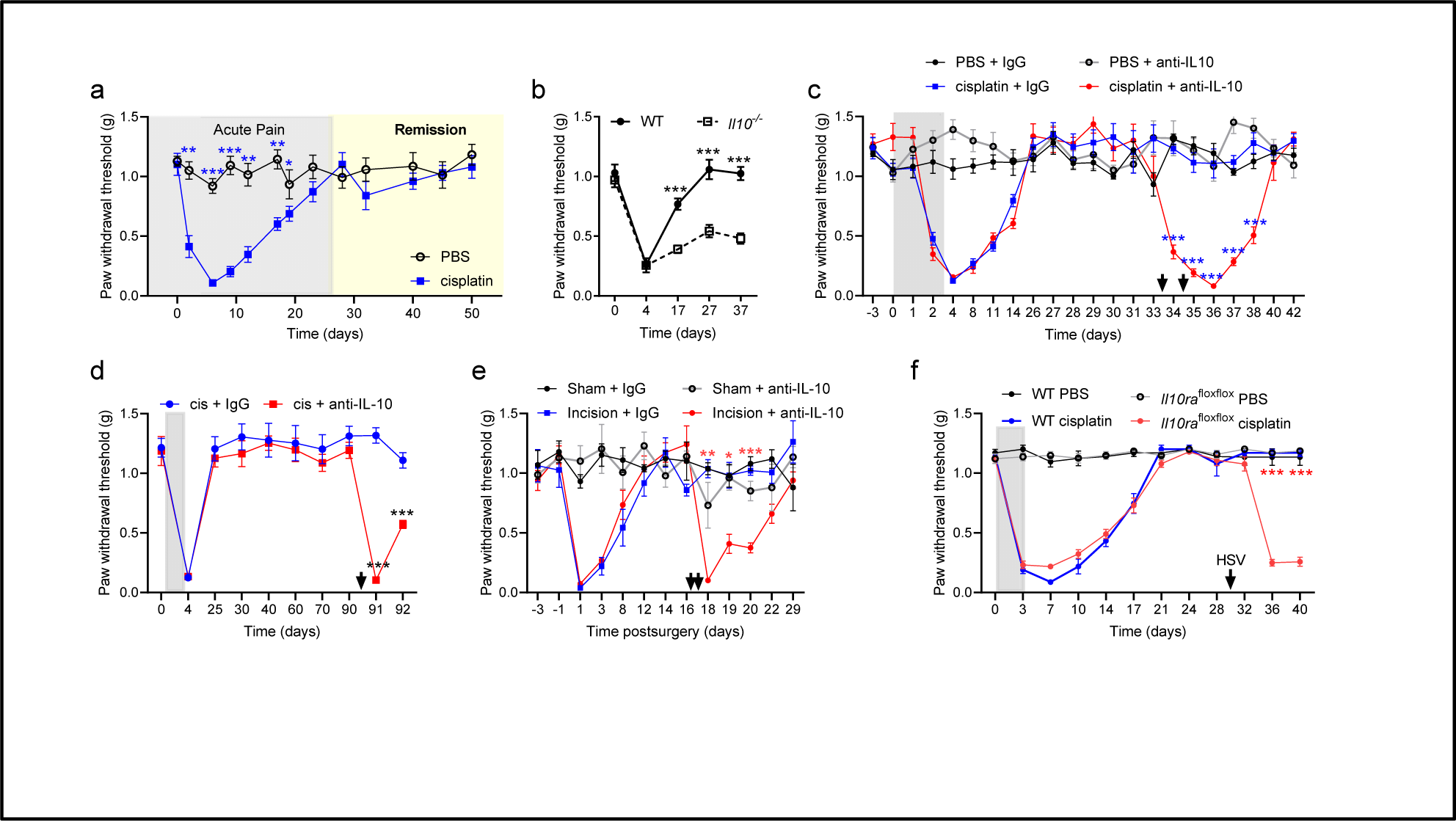
Interleukin-10 signaling is necessary for the maintenance of remission of mechanical pain hypersensitivity. **a**, Mechanical hypersensitivity induced by cisplatin (2 mg/kg/days x 3) peaks at day 8 and reach remission by day 30 (n=6/group). **b**, the lack of IL-10 impairs the resolution of cisplatin-induced mechanical pain hypersensitivity (n=8/group), Two-ways ANOVA genotype effect F(1,14) = 56.0, p<0.0001. **c, d, e**, and **f**: gray area indicate cisplatin/PBS administration and black arrows indicate injection of neutralizing anti-IL-10 (**c-d**) or HSV-mCherry-Cre vector (**f**). **c**, Intrathecal injection of neutralizing anti-IL-10 during remission reinstates mechanical hypersensitivity in cisplatin-treated mice (n=10 mice/group). **d**, Intrathecal injection of neutralizing anti-IL-10 90 days after cisplatin administration reinstates mechanical hypersensitivity (n=10/group. **e**, Mechanical hypersensitivity induced by surgical incision reaches remission by 14 days and intrathecal injection of neutralizing anti-IL-10 on day 16-18 reinstates mechanical hypersensitivity in incision-treated mice (n=4/group). **f**, HSV-mCherry-Cre vector injection on day 31 reinstated mechanical pain hypersensitivity in cisplatin-treated *Il10ra*^floxflox^ mice (n=4-9/group).

Neuropathic pain is also associated with cold hypersensitivity. Thermal place preference was used to assess non-reflexive cold sensitivity/avoidance (**Figure 2A**). Cisplatin induced non-reflexive cold avoidance at day 8 (**Figure 2B**). Similar to von Frey, mice returned to baseline levels on day 32. Inhibition of IL-10 signaling reinstated cold avoidance in cisplatin-treated mice but had no effect on PBS-treated control mice (**Figure 2C**). Virally mediated deletion of IL-10R1 on DRG neurons reinstated cold avoidance in cisplatin-treated mice in remission (**Figure 2D**). In addition, to behavior, a critical feature of cisplatin-induced neuropathic pain is the upregulation of spinal astrocyte marker Glial Fibrillary Acidic Protein (GFAP) (Krukowski et al. 2016, Chiang et al. 2019). GFAP is upregulated in the lumbar spinal cord of cisplatin-treated mice at day 8 when pain hypersensitivity and cold avoidance are present and GFAP levels went back to normal when pain behaviors were in remission (**Figure 2E**). Neutralization of spinal IL-10 reinstated upregulation of GFAP expression in cisplatin-treated mice (**Figure 2E**). These data indicate that IL-10 signaling is essential to maintain remission and prevent relapse to pain.

**Figure 2.**
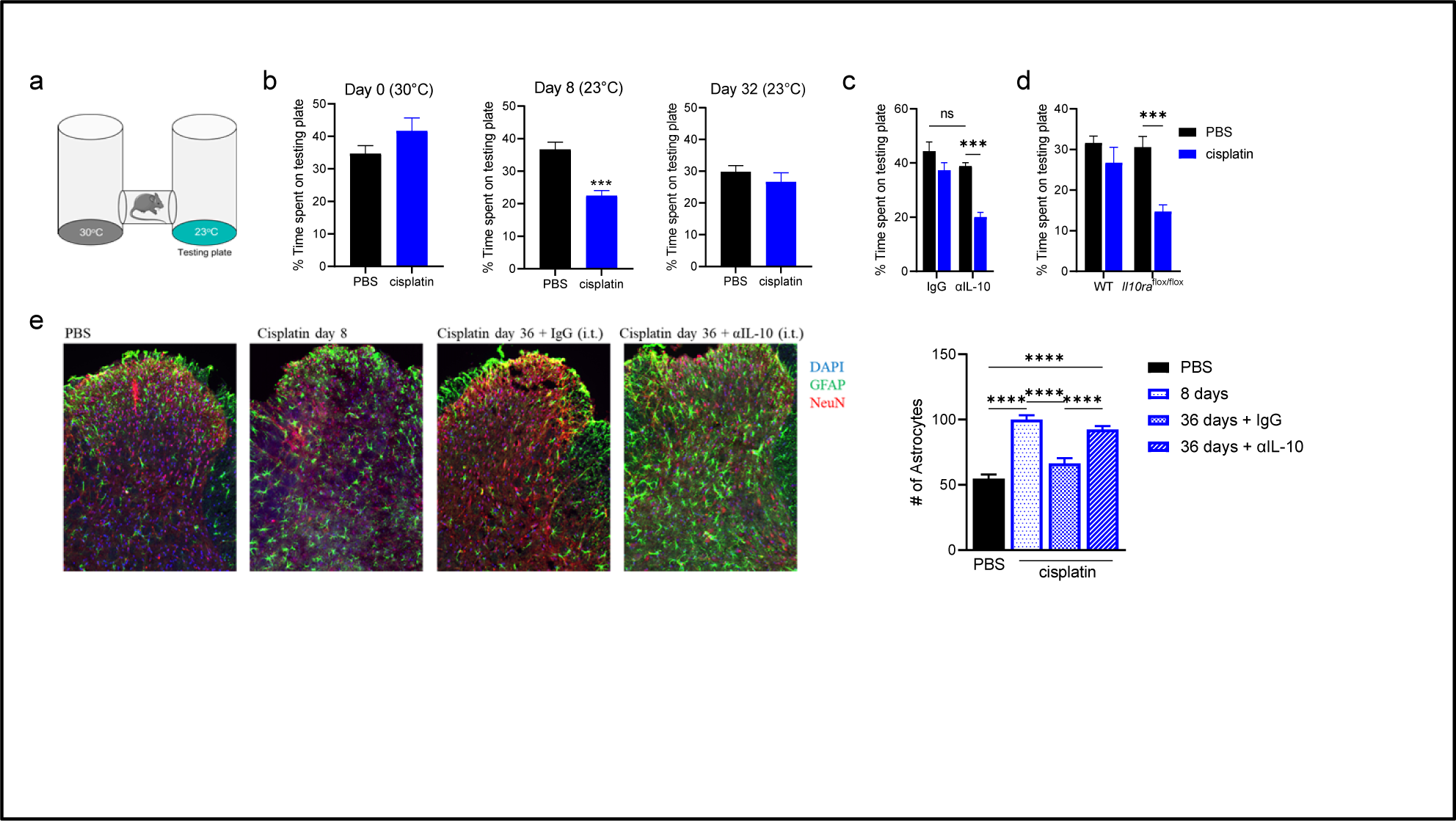
Inhibition of IL-10 signaling during remission reinstated cold pain hypersensitivity and spinal astrogliosis. **a**, Representation of the thermal place preference (TPP). **b**, No preference is shown in the TPP test when both plates are at 30⁰C. Cisplatin-treated mice spend less time on the testing plate on day 8 compared to PBS-treated mice but on day 32, cisplatin- and PBS-treated mice spend equal amount of time on the testing plate (n=6/group). **c**, On day 36, intrathecal injection of neutralizing anti-IL-10 reinstates cold hypersensitivity in cisplatin-treated mice (n=6/group), Two-ways ANOVA cisplatin x antibody interaction F(1,20) = 5.81, p=0.03. **d**, Intraplantar injection of HSV-mCherry-Cre vector reinstates cold hypersensitivity in cisplatin-treated *Il10ra*^floxflox^ mice (n=4-9/group). **e**, The number of GFAP-positive cells (green) in the dorsal horn of the spinal cord increases 8 day after cisplatin and goes back to baseline level on day 36. On day 36, intrathecal injection of neutralizing anti-IL-10 increases the number of GFAP-positive cells (astrogliosis) (n=4-8/group), one-way ANOVA F(3,20) = 36.8, p<0.0001.

### IL-10 is mainly produced by meningeal macrophages

To identify the cellular source of IL-10 in cisplatin- and PBS-treated mice, on day 33-35, when remission of pain was observed, flow cytometry was performed on spinal cord and brain immune cells from IL-10 GFP reporter mice. Unexpectedly, IL-10-producing leukocytes were not detected in the spinal cord nor in the brain (**Figure S2A**). But IL-10-producing leukocytes were present in the spinal meninges (**Figure 3A**). High dimensional data analysis was used to identify meningeal immune cell populations (**Figure 3B, Fig S3A**). IL-10 was produced by different types of leukocytes; GFP+ cells were mostly identified as resident border-associated macrophages (BAMs, CD45^mid^, CD11b+, F4/80+, CD44-) and T cells (**Figure 3C, Fig S3B**). The BAM identify of the IL-10-producing resident macrophages (CD11b+, CD44-) was confirmed by the detection of two additional markers; Lyve1 and CD206 (**Figure S3C**). Recently, CD11b^+^ CD11c^high^ myeloid cells in the spinal cord, were associated with the remission of pain (Kohno et al. 2022). Here, meningeal CD11b+ CD11c+ double positive cells are more important producers of IL-10 than CD11b+ CD11c-cells (**Figure S4**). BAMs in the spinal meninges were identified as the main producer of IL-10. The expression of spinal meningeal IL-10 was increased in cisplatin-treated mice in remission compared to PBS-treated mice (**Figure 3D**).

**Figure 3.**
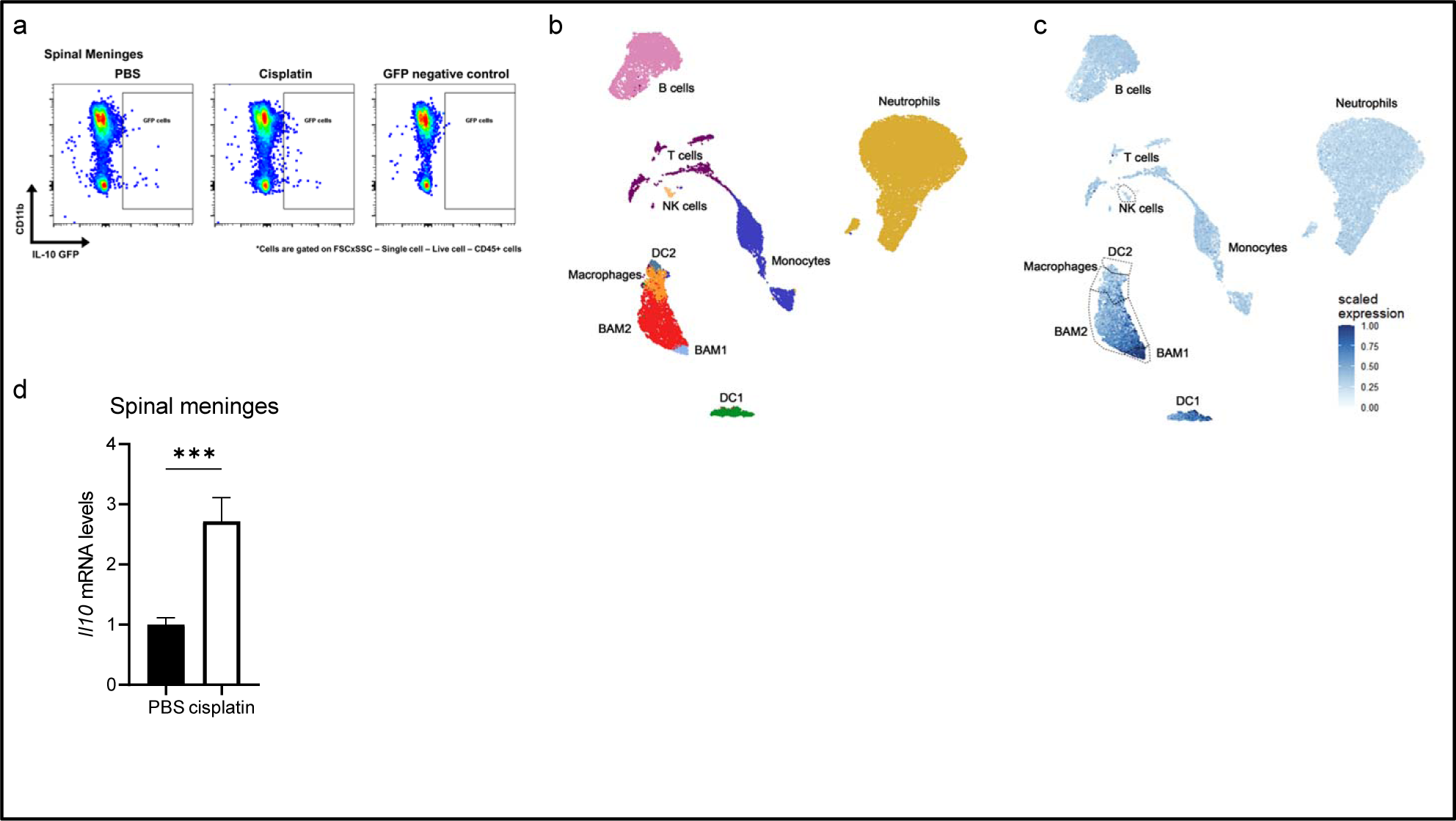
Meningeal macrophages are the cellular source of IL-10. **a**, Representation of CD11b vs. GFP plot from single live CD45+ cell gate isolated from spinal meninges from PBS- and cisplatin-treated IL-10 GFP reporter mice and WT mice (negative control). **b**, UMAP of high dimensional flow cytometry reveals the immune cell composition of the spinal meninges from cisplatin treated mice (n=4). **c**, Identification of IL-10 producing immune cells in the spinal meninges. IL-10 is mainly produced by border-associated macrophages (meningeal resident macrophages). **d**, *Il10* mRNA levels in the spinal meninges increases in cisplatin-treated mice (n=9/group), unpaired t-test df = 16, p=0.0007.

### Inhibition of STAT3 reinstates pain

The main signaling pathways downstream of IL-10R1 are activation of signal transducer and activator of transcription (STAT)-3 (Weber-Nordt et al. 1996, Takeda et al. 1999). Injection of STAT3 inhibitor (1 mg/kg) during remission reinstated mechanical pain hypersensitivity and cold avoidance in cisplatin-treated mice in both sexes (**Figure 4A, B, S1D**). The main effect of STAT3 is the regulation of gene expression. To confirm the genomic effect of STAT-3 inhibition, the expression level of cyclinD1, a well-known target of STAT3 (Bromberg et al. 1999), was assessed (**Figure 4C**). STAT3 DNA binding activity was measured in the DRG during remission. Cisplatin increased STAT3 DNA binding, which is reduced by STAT3 inhibitor and by intrathecal anti-IL-10 antibody (**Figure 4D**). These data show that IL-10 impacts the transcriptional activity of STAT3 during remission.

**Figure 4.**
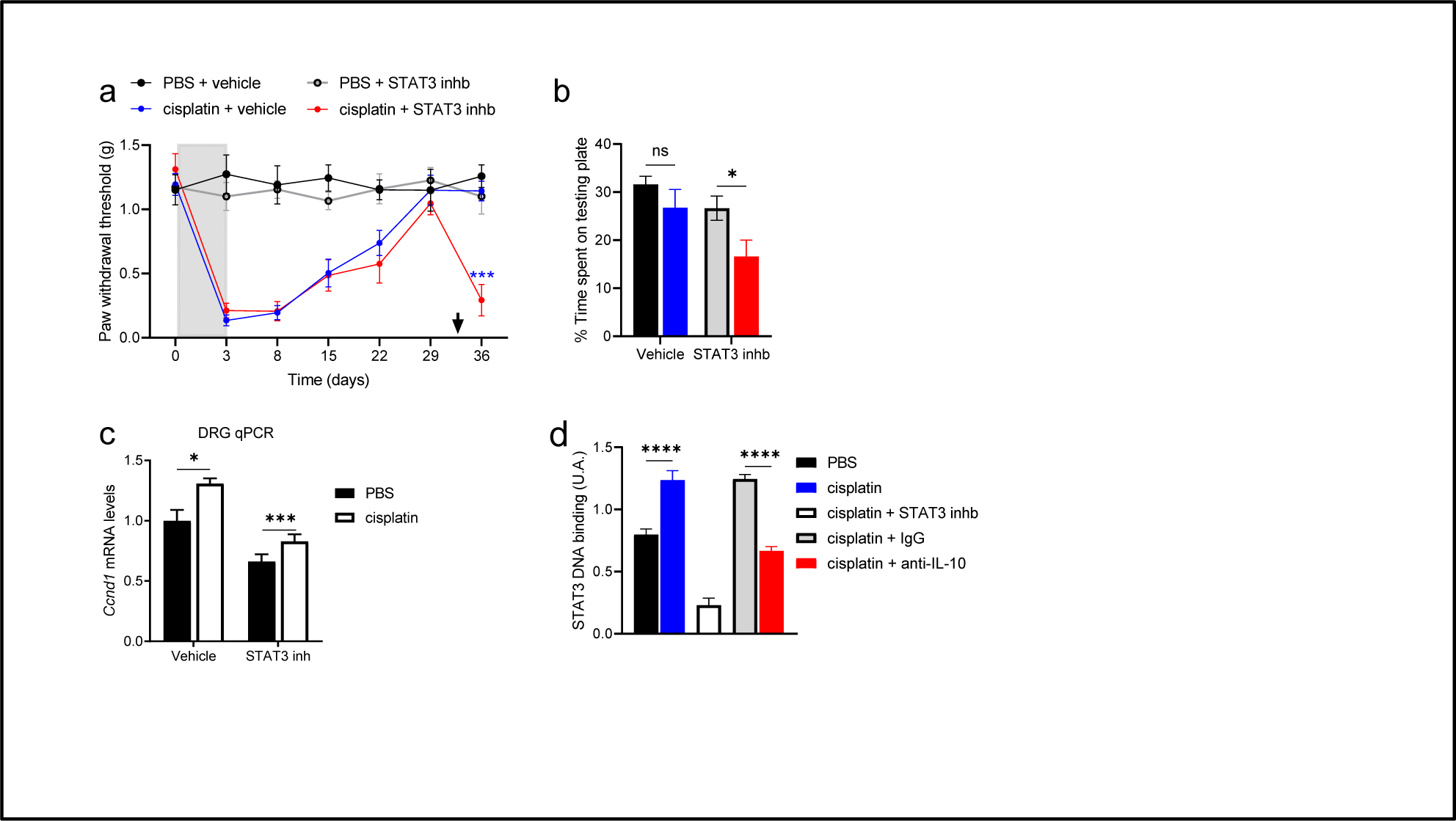
STAT3 contributes to the maintenance of remission from neuropathic pain. **a**, administration of STAT3 inhibitor (1 mg/kg, BP-1-102) on day 30 (arrow) reinstates mechanical hypersensitivity in cisplatin-treated mice (n=8/group). Gray area indicates PBS or cisplatin treatment (2 mg/kg/day x3). **b**, injection of STAT3 inhibitor reinstates cold hypersensitivity assessed by TPP (n=4-6/group), two-ways ANOVA stat3 effect F(1,16) = 5.89, p=0.03. **c**, *Ccnd1* mRNA level in DRG is regulated by STAT3 (positive control) (n=5/group). **d**, On day 36, cisplatin increases the binding of STAT3 to DNA, the binding is blocked by STAT3 inhibitor (positive control) and intrathecal injection of neutralizing anti-IL-10 reduces STAT3 binding to DNA (n=6/group), one way ANOVA F(4, 25) = 68.8, p<0.0001.

### IL-10 signaling upregulates **δ**OR during remission

STAT3 is a transcriptional factor that mediates IL-10 effects mostly by regulation of gene expression (Benkhart et al. 2000, Ding et al. 2001). Moreover, chronic pain has been associated with transcriptomic reprogramming in DRG neurons (Laumet et al. 2015, North et al. 2019). These observations suggest that changes in gene expression in the DRG are critical to the regulation of chronic pain. To determine the mechanisms underlying the maintenance of remission of pain by IL-10 signaling in the DRG, following remission and relapse induced by anti-IL-10, an RNA sequencing was performed on DRG samples on day 36 (**Figure S5A and supplementary data)**. Principal component analysis showed that the samples clustered by treatments (**Figure S5B**). Overall, 109 genes were differentially regulated in “Remission” and 153 genes in “Relapse” (**Figure 5A**). Only *Jspr1* and *Oprd1* were significantly differentially expressed in both conditions (**Figure S5C**). The top 10 genes differentially expressed with the lowest p-value for “Remission” (PBS vs. cisplatin) and “Relapse” (cisplatin + IgG vs. cisplatin + anti-IL-10) are presented in **Supplementary Table 1**. Interestingly, *Socs3* a signaling molecule downstream of IL-10R1 is among these genes.

**Figure 5.**
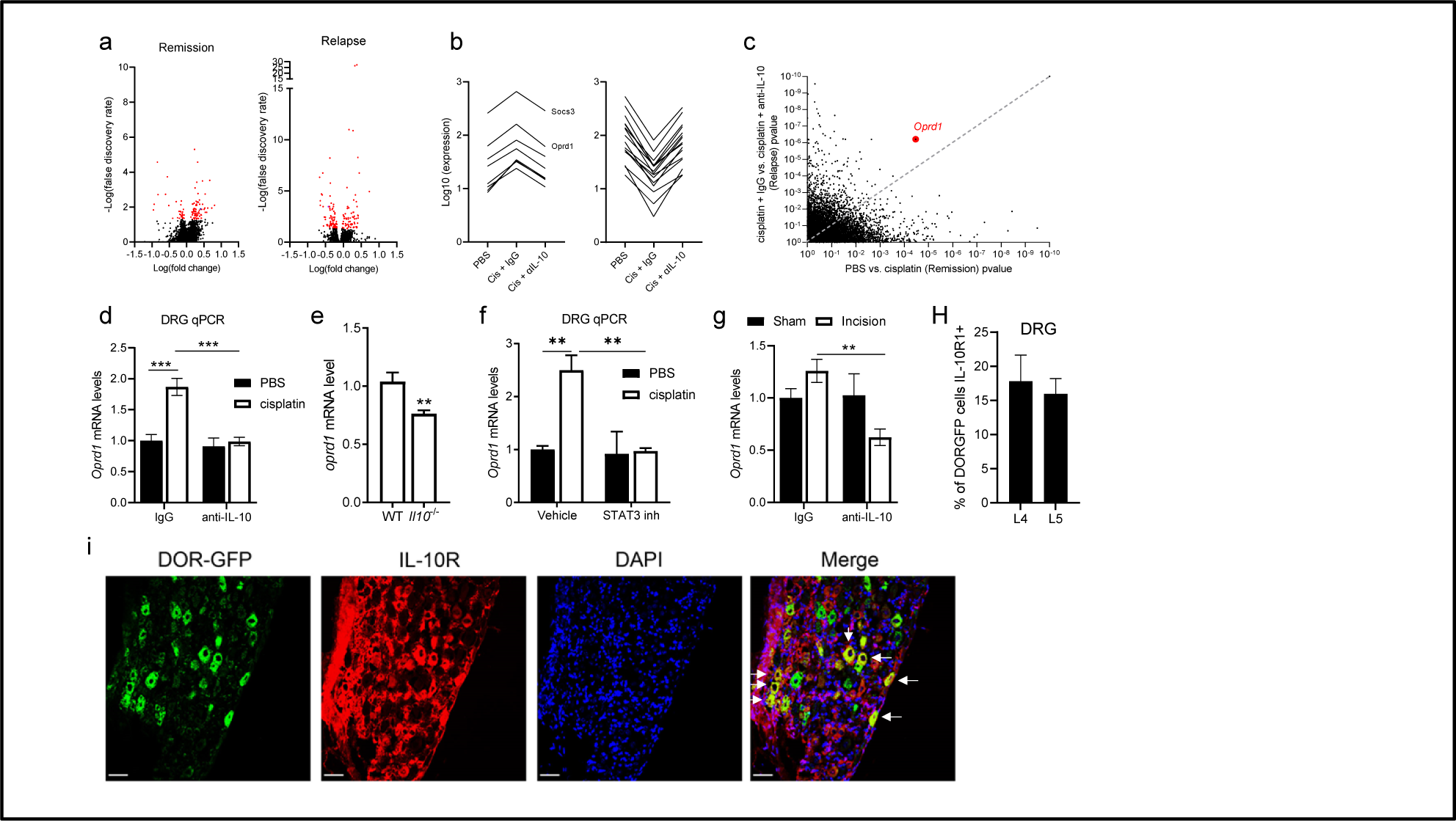
During remission DRG *Oprd1* is upregulated by IL-10 signaling. **a**, Volcano plot of RNA sequencing from lumbar DRGs on day 36 comparing PBS-vs. cisplatin treated mice “Remission” and cisplatin+IgG vs. cisplatin+αIL-10 “Relapse”. Red plots indicate statistical significance difference for genes with –log(false discovery rate) >1.3 (n=4/group). **b**, Line plots of genes showing a normalization of their expression in relapsing mice (RemissionFC>2 and RelapseFC<0.5), grouped according to their expression change in the remission group: upregulated (n=8 genes) and downregulated (n=16 genes). **c**, Comparison of p-values of genes differentially expressed in Remission and Relapse, the gene with the lowest p value in both conditions is Oprd1 (Remission p= 3.38×10^-5^, Relapse p=6.13×10^-7^). **d-g**, qPCRs showing that *Oprd1* mRNA levels is increased in cisplatin-treated mice in remission and this upregulation is blocked by intrathecal neutralizing αIL-10 (n=11-12/group) two-way ANOVA cisplatin x αIL-10 interaction F(1,42) = 11.9, p=0.0013 (**d**), *Il10*^-/-^ (n=8/group) unpaired t-test df=14, p=0.006 (**e**), STAT3 inhibitor (n=5/group) two-way ANOVA cisplatin x STAT3 inh interaction F(1,16) = 7.96, p=0.01 (**f**), and in remission from postoperative pain (n=6-7/group) two-way ANOVA surgery x αIL-10 interaction F(1,22) = 6.77, p=0.02. **h**, quantification of cells that co-express δOR and IL-10R1 in DRG section from DOR-GFP reporter mice (n= 4 mice). **i**, Representation of DRG section from DOR-GFP mice; δOR (green), IL-10R1 (red), and DAPI (blue), white arrow indicate cells that are co-labeled for green and red.

Gene(s) important for the balance between remission and relapse were identified as differentially expressed during remission, but their expression was normalized during relapse: RemFC>2 and RelapseFC<0.5 (*Socs3, Oprd1, Ston2, Palm2*, Gm26724, D030047H1, *Tas1r3, Fam205c*) or RemFC<0.5 and Relapse>2 (*Gfap, Itih3, Ckmt2, Ckm, Myh4, Myl1, Tnnt3, Atp2a1, Trdn, Myoz1, Cox6a2, Mylpf, Acta1, Tnni2, Tcap, Myh1*) (**Figure 5B**). Among these 25 genes, only *Oprd1* expression was significantly different in both conditions (**Supplementary Table 2**). Then, all differentially expressed genes from “Remission” and “Relapse” conditions are compared, the gene with the lowest p-value in both conditions is *Oprd1*, a gene encoding δ-opioid receptor (δOR) (**Figure 5C**). The upregulation of *Oprd1* in the DRG during remission was confirmed in a different cohort of mice by qPCR and this increase was blocked by anti-IL-10 (**Figure 5D**) or in *Il10*^-/-^ mice (**Figure 5E**). No change was observed in the spinal cord for *Oprd1 nor Oprm1* (encoding the morphine receptor µOR) (**Figure S6A**). The expression of *Oprm1* in the DRG was not affected by cisplatin nor IL-10 signaling (**Figure S6B,C**). Inhibition of STAT3 also blocked the upregulation of *Oprd1* (**Figure 5F**). Similarly, in the postoperative model, inhibition of IL-10 signaling reduced *Oprd1* expression during remission (**Figure 5G**). Treatment of DRG cell line with recombinant IL-10 and cisplatin upregulated the expression of *Oprd1* (**Figure S6D**). Immunostaining of IL-10R1 in DRG section from δOR-GFP mice showed that IL-10R1 and δOR are co-express in ∼15-18% of DRG neurons (**Figure 5H, I**), which is consistent with mouse single cell-RNA-sequencing (scRNA-seq) data (**Figure S6E**)(Zeisel et al. 2018).

### δOR in DRG critically controls the remission of pain

Consistently with the upregulation of *Oprd1* observed here, previous studies showed that systemic injection of δOR antagonist reinstates pain hypersensitivity in models of chemotherapy-induced neuropathic pain, postoperative pain, and inflammatory pain (Marvizon et al. 2015, Inyang et al. 2021). To take advantage of genetic specificity and determine if the contribution of δOR is taking place at the level of DRG neurons, HSV-Cre viral vector was injected in *Oprd1*^floxflox^ mice. The downregulation of *Oprd1* in the DRG was confirmed by qPCR and mCherry expression (**Figure S7A-C**). Genetic downregulation of *Oprd1* in DRG neurons reinstated pain hypersensitivity and cold avoidance in cisplatin-treated mice in both sexes (**Figure 6A, B**). Interestingly, deletion of *Oprd1* in DRG neurons during remission is sufficient to re-instate astrogliosis in the dorsal horn (**Figure 6C, D**). Then, the hypothesis that over-expression of δOR compensates for the lack of IL-10 signaling was tested. *Il10ra*^floxflox^ mice were treated with cisplatin and once they reached remission injected with HSV-Cre which triggered relapse to pain. Following relapse, these mice were injected with HSV-mCherry-Oprd1 to cause an overexpression of δOR. Overexpression of δOR triggered resolution of mechanical pain hypersensitivity in cisplatin-treated Il10ra^floxflox^ mice (**Figure 6E**) but failed to reverse cold avoidance (**Figure 6F**) following HSV-Cre treatment.

**Figure 6.**
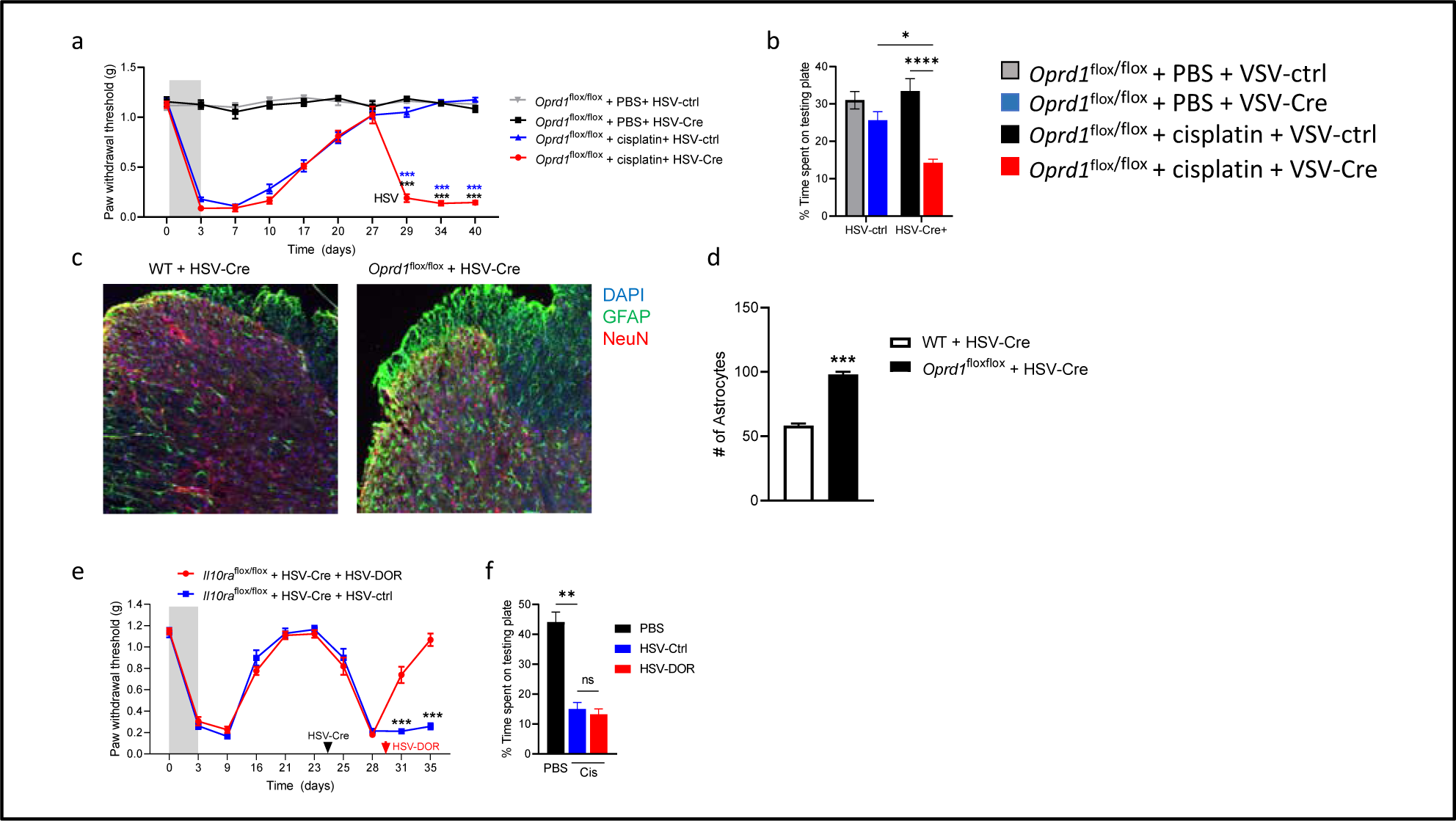
Inhibition of δ-Opioid receptor signaling in the DRG triggers relapse to pain. Intraplantar injection of HSV-mCherry-Cre vector on day 28 reinstated mechanical (**a**) and cold pain hypersensitivity (**b**) in cisplatin-treated *Oprd1*^loxflox^ mice (n=7-9/group). **c**, Representative image of GFAP staining in the dorsal horn. **d**, the number of GFAP-positive cells is increased by the deletion of *Oprd1* in the DRG neurons (n=4/group) one-way ANOVA F(4,17) = 51.8, <0.0001. **e**,. Intraplantar injection of HSV-mCherry-Cre vector on day 23 reinstates mechanical hypersensitivity in cisplatin-treated *Il10ra*^floxflox^ mice and subsequent injection of HSV-mOprd1 vector restores mechanical pain hypersensitivity (**e**) but not cold hypersensitivity (n=8-9/group) (**f**). **a, e**, gray area indicates cisplatin treatment (2 mg/kg/day x3).

### IL-10 signaling regulates **δ**OR analgesic activity during remission

Corder et al., showed that µOR agonist enhanced µOR-mediated analgesia in the hot plate during remission (Corder et al. 2013). During remission, mice were treated with saline, SNC80 (δOR specific agonist) and DAMGO (µOR agonist) before placed on the hot plate. SNC80 did not affect the latency to react to noxious heat, but the analgesic effect of DAMGO was increased during remission consistently with previous report (Scherrer et al. 2009, Corder et al. 2013) (**Figure S7D**). To test whether non-nociceptive δOR-mediated behaviors are affected by IL-10 signaling, SNC80-induced enhanced locomotor activity (LMA) was monitored (Longoni et al. 1998, Spina et al. 1998). As expected SNC80 induced increased LMA in WT and in *Il10*^-/-^ mice (**Figure S7E**) indicating that IL-10 signaling does not regulate δOR linked enhanced locomotion. HSV-Cre-induced deletion of *Oprd1* blocked the analgesic effects of SNC80 confirming the importance of DRG δOR (**Figure S8**).

δOR in DRGs in usually inactive in naïve animals and get activated upon injury (Brackley et al. 2016, François et al. 2018). As δOR in DRGs plays a critical role in remission, it is important to test whether δOR analgesic activity is also enhanced during remission of pain. Following remission from pain hypersensitivity, mice were treated with SNC80 (10 mg/kg). Injection of SNC80 increased the pain sensitivity threshold in cisplatin-treated mice but had no effect on PBS-treated mice. Neutralization of IL-10 prevented the analgesic effect of SNC80 (**Figure 7A**). Similar data were observed after intrathecal injection (**Figure 7B**). SNC80 administered either i.p. or i.t. failed to induce analgesia in Il10-/-mice (**Figure 7C**). SNC80 analgesic effects were also absent in *Avil*^Cre^-*Il10ra* mice during remission (**Figure 7D-F**) and at the peak of acute pain hypersensitivity, 7 d post cisplatin injection (**Figure S7F**. In *Il10ra*^floxflox^ mice, when relapse was induced by injection of HSV-mCherry-Cre to remove IL-10R1 on DRG neurons, injection of SNC80 failed to alleviate pain hypersensitivity (**Figure 7G**). Overall δOR agonism enhanced pain sensitivity in mice in remission with functional IL-10 signaling in the DRG.

**Figure 7.**
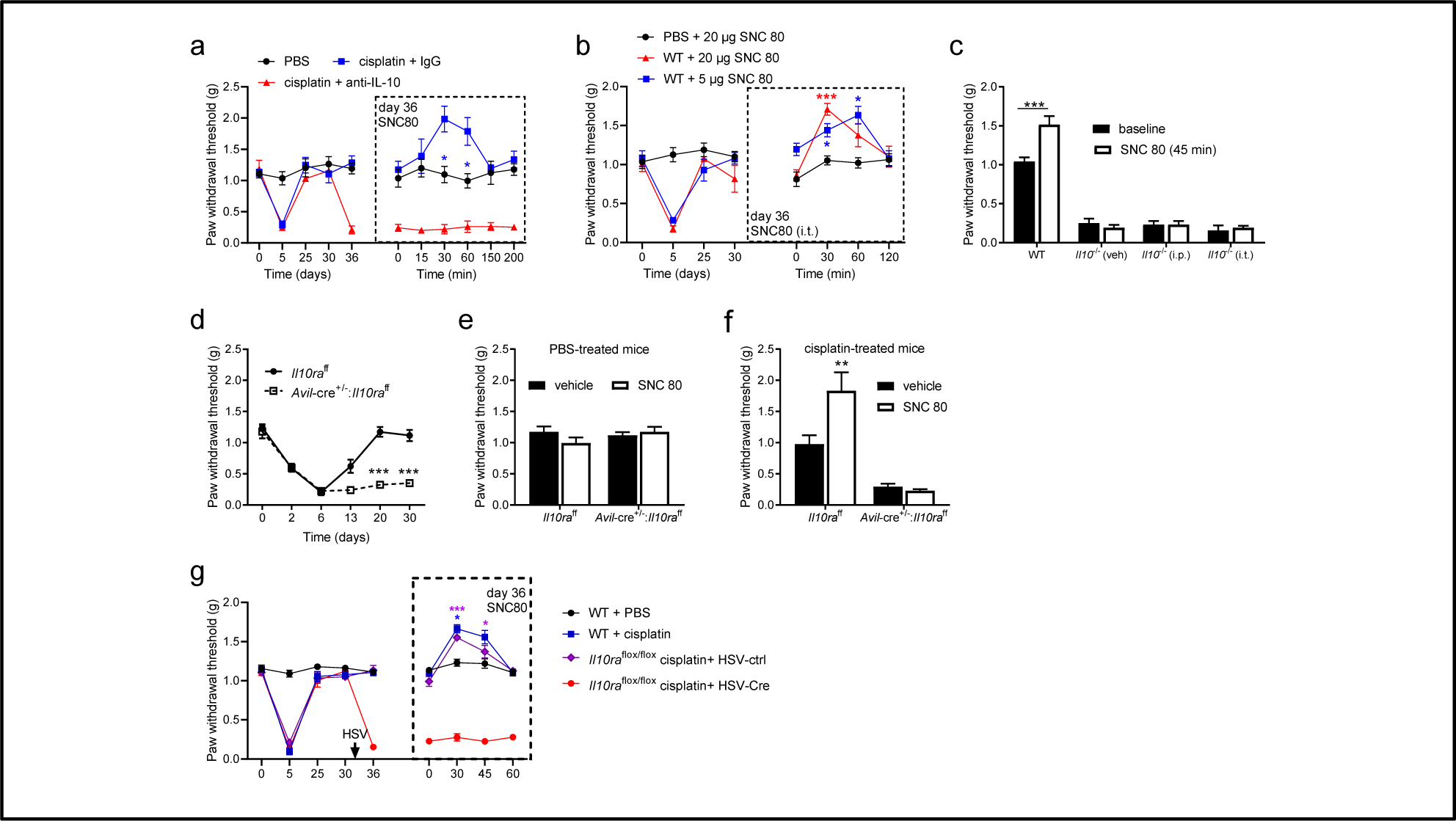
δ-Opioid receptor selective agonist induces analgesia during remission in mice with intact IL-10 signaling. **a**, Mechanical sensitivity of mice treated with PBS, cisplatin and αIL-10 (intrathecal injection day 33-34), on day 36 mice are treated with SNC 80 (10 mg/kg) and mechanical sensitivity is recorded for 200 min; activation of δOR induces analgesia in cisplatin-treated mice with intact IL-10 signaling (n=6/group). **b**, intrathecal injection of SNC 80 induces analgesia in cisplatin-treated mice (n=6/group). **c**, On day 30 after cisplatin administration, mice received SNC 80 either i.p. 10 mg/kg or intrathecal (20 µg) and 45 min later assessed for mechanical sensitivity; the lack of IL-10 impairs the analgesic effect of SNC80 (n=6/group). **d**, Absence of IL-10 signaling in DRG neurons prevents mice to reach remission (n=6/group). **e,f**, On day 32, mice received ip injection of SNC80 and mechanical sensitivity was assessed 45 min after; δOR agonism induces analgesia in cisplatin-treated *Il10ra*^floxflox^ mice but not in mice that lack IL-10R1 on advillin-positive neurons (same mice as **d**). **g**, δOR agonism induces analgesia in cisplatin-treated mice in remission but not in mice that lack IL-10R1 on DRG neurons (n=5-9/group).

## Discussion

These data strongly suggest a new concept that defines remission as an active regulatory event involving neuroimmune interactions to suppress a sensitized state. Our novel findings indicate that IL-10 remains persistently upregulated, is critical to upregulate δOR analgesic activity to maintain remission and prevent relapse to pain. These data are observed in two models with different etiology: chemotherapy and plantar incision. Our study also demonstrated that IL-10 is mainly produced by border-associated macrophages (BAMs) in the spinal meninges. Despite the sexual dimorphism of the immune system and pain prevalence, here behavioral and molecular data were similar in male and female mice which is consistent with other studies on IL-10 signaling (Laumet et al. 2020, Prado et al. 2021, Guo et al. 2022).

The present work expands the concept that the somatosensory/nociceptive system remains sensitized while there is no sign of pain-like behaviors anymore (Campillo et al. 2011, Corder et al. 2013, Melemedjian et al. 2014, Taylor et al. 2019). The role of the opioid system has been clearly established in remission of pain (Campillo et al. 2011, Taylor et al. 2014, Marvizon et al. 2015, Inyang et al. 2021) and recently a role of neuroimmune interactions has been reported as well (Kohno et al. 2022). However, our study is the first to show the contribution of IL-10 and to link neuroimmune interactions and the opioid system in remission from pain. Some of the mechanistic differences between the different findings on remission (Campillo et al. 2011, Marvizon et al. 2015, Basu et al. 2021) are likely to result from different etiology specific to each pain model. For example, analgesic effect of δOR is more associated with neuropathic pain models (Cahill et al. 2022), while µOR is predominant to alleviate inflammatory pain (Taylor et al. 2014, Gendron et al. 2016, Kandasamy et al. 2021). The fact that IL-10, like µOR (Corder et al. 2013), is necessary for the initial resolution of pain and to prevent relapse suggest a continuum in the mechanisms underlying resolution and maintenance of remission. Although this study was restricted to the DRG and spinal cord, we cannot rule out a role for the descending inhibitory pathways (Chen et al. 2014). A potential explanation for the persistent upregulation of IL-10 is ongoing DRG neuron activity, as C-fiber activity stimulates IL-10 expression (McKelvey et al. 2015). As IL-10 suppresses DRG neuron hyperactivity (Laumet, 2020; Guo, 2022), it might act as negative feedback for preventing neuronal hyperactivity.

IL-10 levels have been associated with relief and remission from chronic pain (Uçeyler et al. 2007, Glocker et al. 2011, Zorina-Lichtenwalter et al. 2018, Banimostafavi et al. 2021, Bussmann et al. 2022). Additionally, in preclinical models, the analgesic role of IL-10 has been clearly established (Milligan et al. 2006, Sloane et al. 2009, Milligan et al. 2012, Grace et al. 2017, Vanderwall et al. 2018). Here, we critically identified the cellular source of IL-10 by taking advantage of GFP reporter mice and high-dimensional spectral flow cytometry. We found that IL-10 is mostly produced by BAMs in the spinal meninges. In line with these findings, BAMs were found to be essential for the resolution of sham-surgery (Niehaus et al. 2021). CD11b+ CD11c^high^ microglia were recently proposed to maintain remission (Kohno et al. 2022). Interesting, in this article some CD11b+ CD11c+ cells are also outside of the spinal cord parenchyma and probably in the spinal meninges. Here CD11+ CD11c+ BAMs are an important producer of IL-10, therefore we made the suggestion that IL-10+ CD11b+ CD11c+ BAMs might be a common cell type in both studies and be essential for the regulation of remission and relapse. In addition to macrophages, T cells also produced IL-10, which is consistent with previous studies (Krukowski et al. 2016, Laumet et al. 2019, Laumet et al. 2019, Durante et al. 2021). We acknowledge that our harvested spinal meninges contain dorsal root leptomeninges. We cannot exclude that IL-10+ BAMs are in the dorsal root leptomeninges which is a direct route to DRG neurons (Maganin et al. 2022). The strong expression of IL-10R1 in DRG (Laumet et al. 2020, Prado et al. 2021) reinforces this hypothesis.

Although, the analgesic effect of IL-10 was thought to be only mediated by immunosuppression (Sabat et al. 2010), We and others have shown a direct analgesic role of IL-10 on neurons (Laumet et al. 2020, Prado et al. 2021, Guo et al. 2022). We expanded our previous work to reveal a critical role of IL-10 in regulating DRG transcriptome and δOR more specifically. IL-10R1 and δOR are expressed by the same cells in the DRG. This result is in line with the scRNAseq data.

We identified Oprd1 as a new target of IL-10/STAT3 axis in DRG. Our data show that δOR remains inactive in DRG of naïve mice but gets activated during remission. This observation is consistent with previous report that show that injury/inflammation is necessary to activate the analgesic effect of δOR in DRG to counterbalance the development of pain hypersensitivity (Kabli et al. 2007, Obara et al. 2007, Gaveriaux-Ruff et al. 2011, Pradhan et al. 2013, Bardoni et al. 2014, Brackley et al. 2016, Gonçalves et al. 2021). The upregulation of IL-10 which is often associated with injury or inflammation may explain the activation of δOR in these models. Remission results from a constant suppression of pain hypersensitivity by opioid signaling. This tonic suppression is necessary to restrain spinal astrogliosis. The molecular mechanism triggering the activation of δOR in DRG requires further investigations. On the other hand, overexpression of δOR in DRG partially bypassed the lack of IL-10 signaling on neurons. However, this overexpression of δOR did not have the anticipated effect on cold allodynia suggesting that δOR play a limited role in thermal hypersensitivity (Wang et al. 2018).

Taken together, we demonstrated novel neuroimmune-opioid crosstalk underlying the balance between remission and relapse of pain. Based on these findings, we propose that IL-10 signaling might be both a biomarker for the risk of relapse and a therapeutic target for relapsing chronic pain.

## Method

### Animals

All animal experiments were approved by the Institution Animal Care and Use Committee (IACUC) at Michigan State University, The University of Texas MD Anderson Cancer Center, and University of North Carolina. Mice were housed and bred in university animal care facilities prior to experiments and treatment. Animals had *ad libitum* access to food and water and were on a 12 hr non-inverted light/dark cycle. Mice were randomized to treatment groups. The following mouse strains were purchased from Jackson Laboratories (Bar Harbor, ME) and bred before use: C57Bl6 (WT, JAX#000664); *Avil*^Cre^ (Zhou et al., 2010) (JAX#032536), and *Il10ra*^fl/fl^ (Liu et al., 2012) (JAX#028146) to generate *Avil*^Cre^:*Il10ra^fl/fl^*(Laumet et al. 2020); IL-10 GFP (JAX#014530) (Madan et al. 2009); *Oprd1fl* (JAX#030075) (Gaveriaux-Ruff et al. 2011); *Il10*^-/-^ (JAX#002251) (Kühn et al. 1993); δOR-GFP (JAX#029012) (Scherrer et al. 2006).

*Models of pain remission*. Transient neuropathic pain followed by remission was induced by intraperitoneal injection (i.p.) of cisplatin (Cat# 1134357; Sigma Aldrich; St. Louis, MO) 2 mg/kg x 3 days (Laumet et al. 2020, Inyang et al. 2021). Postoperative pain was induced by making a 5-mm longitudinal incision with a number 11 scalpel blade (Cat# 12-000-133; Fisher Scientific) in the skin of the left hindpaw and the underlying muscle tissue 2 mm below the heel. The wound was closed using a 5-0 suture (Cat# S-G518R13; AD Surgical; Sunnyvale, CA), followed by a 200-µL subcutaneous injection of gentamicin (5 mg/kg; Cat# G1272; Sigma Aldrich) (Inyang et al. 2019, Inyang et al. 2021).

Animals were terminated by CO_2_ exposure and perfused with ice-cold PBS. Tissues for qPCR and ELISA were snap frozen. Tissues for immunohistochemistry were incubated in ice-cold 4% paraformaldehyde and cryoprotected with 30% sucrose (Cat# S5-500; Fisher Scientific).

#### Behavior

Mechanical Pain hypersensitivity was measured with von Frey filaments (Cat# BIO-VF-M; Bioseb; Pinellas Park, FL) using the Dixon method. Briefly, the experiment begins by testing the response to a 0.4g filament. If there is no response, the next filament with a higher force is tested; if there is a response, the next lower force filament is tested. This continues until 5 readings are obtained after the first change of direction, and the sequence of outcomes is recorded (Chaplan et al. 1994) as in previous work (Laumet et al. 2020, Inyang et al. 2021).

Cold avoidance was measured using the thermal place preference apparatus (Cat# 35250; Ugo Basile S.R.L; Gemonio VA, Italy). Two enclosed hotplates were connected by a small, narrow tunnel. Animals were first habituated to the plates 24 hours prior to testing. The mice are then returned to the apparatus with both plates set to 30 C and recorded for 10 min. One of the hotplates is then set to 23 C and the mice are returned to the apparatus and recorded for an additional 10 min. These recordings were analyzed using the Toxtrack software(https://toxtrac.sourceforge.io) (Rodriguez et al., 2018).

Response to noxious heat was measured by hotplate (Cat# 35300; Ugo Basile). Animals were first habituated to the device at 30 C. Then, animals were placed individually on the plate at 52 C for a maximal period of 1 min. Latency to withdraw a hind paw was recorded (Cancer Exo?).

To record locomotor activity (LMA), mice were placed in new empty cage for 5 min and videotaped from above. Their activity was analyzed with Toxtrac software.

All behaviors were performed by experimenters blinded to the experimental conditions.

#### Viral vector

Replication incompetent herpes-simplex virus (HSV) were obtained from Rachael Neve, director of the Gene Delivery Technology Core (GDTC) at Massachusetts General Hospital. Five µl of viral vector (10^9^ infectious units /dilution 1:7 in PBS+10% sucrose + 25 mM HEPES) were injected subcutaneously into both hind paws. The different HSV vectors are described in **Table 3**.

#### Drugs

SNC80 (10mg/kg; Cat# S2812; Sigma-Aldrich); Anti-IL-10 (10 µg; Cat# 4156-50; VWR International; Radnor, PA); [D-Ala^2^, N-Me-Phe^4^, Gly^5^-ol]-Enkephalin acetate salt (DAMGO; Cat# E7384; Sigma); STAT3 inhibitor XVIII, BP-1-102 (1 mg/kg; Cat# 573132; EMD Millipore Corp; Darmstadt, Germany)

### Molecular biology

#### RNA isolation and qPCR

Spinal cord, dorsal root ganglia, spinal meninges samples were collected, snap frozen and stored at −80C. Total RNA was extracted using a modified version of Trizol-chloroform method to improve the quality of the isolated RNA and quantified with high-performing Qubit 4 Fluorometer instrument (Qubit 4, Invitrogen, Carlsbad, CA). A total of 0.5 μg RNA was reverse transcribed into complementary DNA (cDNA) and quantitative real-time polymerase chain reaction was performed using a CFX96 (Biorad). Expression levels were calculated using the ΔΔCT method and normalized to Gapdh. Data are presented as the average relative fold change normalized to the control group. Gene expression was assessed using validated Primetime primers (Integrated DNA technology, Newark, NJ): *Gapdh* (Ex2-3#Mm.PT.39a.1(1)), *Il10 (Ex3-5#Mm.PT.58.13531087(1))*, *Oprd1 (Ex2-3#Mm.PT.58.30543908)*, *Ccnd1 (Ex4-5#Mm.PT.58.28503826)*.

#### Enzyme-linked immunosorbent assay (ELISA)

Serum IL-10 was quantified by ELISA. Blood was collected by cardiac puncture. Samples were allowed to clot at room temperature for 2 hours followed by a centrifuging for 20 minutes at 2000x g following instructions provided from the Quantikine® HS ELISA kit (Cat# M1000B, R&D systems, Minneapolis, MN). 50µL of Assay Diluent RD1W was added to each well followed by 50 μL of standard, control, or sample per well. The plate was covered with the adhesive strip provided and incubated for 2 hours at room temperature on a horizontal orbital microplate shaker (0.12” orbit) set at 500 rpm. The wells were washed 4 times with the Wash Buffer and all of the liquid was removed following the final wash. 100 µL of Mouse IL-10 conjugate was added to each well, the plate was covered and incubated for 2 hours on the shaker. The plate was again washed and aspirated 4 times followed by a 30 minute incubation with 100 μL per well of Substrate Solution in the dark. After 30 minutes, 100 µL of the Stop Solution was added to each well. The optical density of each well was measured within 30 minutes using a microplate reader set to 450 nm.

#### STAT3 DNA binding

Reagents were added to wells as indicated in Table 1 product insert and incubate one hour at room temperature on an orbital shaker. Each well was washed five times with Wash Buffer and 100 μL of Transcription Factor STAT3 Primary Antibody (1:100) per well was added. Samples were incubated for one hour at room temperature on an orbital shaker then washed five times with 200 μL of 1X Wash Buffer. 100 μL of Transcription Factor Goat Anti-Mouse HRP Conjugate (1:100) was added to each well and incubated for an hour at room temperature on the orbital shaker. Samples were again washed five times with 200 μL of Wash Buffer then 100 μL of Transcription Factor Developing Solution was added to each well. Samples were incubated for 30 minutes at room temperature on an orbital shaker while being protected from light. At the end of the 30 min, 100 μL of Transcription Factor Stop Solution was added per well to stop the reaction. Absorbance was read at 450 nm.

**Table 1.** Immunostaining Antibody table.

| Antibody | Manufacturer | Catalog# | Concentration |
| --- | --- | --- | --- |
| Dapi | Thermo Scientific | 62247 | 1:50,000 |
| GFAP (rabbit) | Proteintech | 16825-1 | 1:1000 |
| GFP (goat) | Abcam | ab6673 | 1:2000 |
| Goat anti-Mouse IgG<br>Alexa Fluor 568 | Invitrogen | A-11004 | 1:1000 |
| Goat anti-Rabbit IgG<br>Alexa Fluor 488 | Invitrogen | A32731 | 1:1000 |
| IL10R1 (mouse) | Santa Cruz<br>Biotech | sc-28371 | 1:200 |
| mCherry (rabbit) | Abcam | ab167453 | 1:1000 |
| NeuN (mouse) | Proteintech | 66836-1 | 1:500 |

#### Immunostaining

DRGs were removed, placed in 4% paraformaldehyde overnight, transferred to 30% sucrose for cryoprotection for 24 hrs, then mounted in Optimal Cutting Temperature (OCT; Cat# 23-730-571; Fisher Scientific) compound. DRG sections were cut into 20 μm slices using a cryostat and mounted onto positively charged (Superfrost plus; Cat# 12-550-15; Fisher) slides for immunohistochemistry. Following 3 5-minute washes in 1X phosphate buffered saline (PBS), the slides were then put into a permeabilization solution containing 10% normal goat serum (NGS; Cat# S13150H; R&D Systems, Minneapolis, MN) and 0.2% Triton X 100 (Cat# 9002-93-1; Sigma) in PBS for 30 minutes. This was followed by another series of 5-minute washes in PBS and 1 hr in a blocking solution containing 10% NGS and 0.01% Na azide (Cat# 18-613-272; Fisher) in PBS. Following another PBS wash, the slides were incubated overnight in a primary antibody solution made from the blocking solution (**Table 1**). The next day, the slides were washed again in PBS then incubated in a secondary antibody solution also made from the blocking solution for 1 hr. Following a PBS wash, counterstaining with DAPI, a PBS wash, and a wash in deionized H_2_O, Prolong Gold mounting media was used to mount coverslips.

#### RNA-sequencing

Whole-genome RNA sequencing was used to identify transcriptional changes induced by cisplatin in the DRG. Total RNA was isolated on day 36 (**Fig. S5A**) with the RNeasy MinElute Cleanup Kit (Qiagen, Hilden, Germany). Libraries were prepared with the Eukaryotic RNA-seq (20M raw reads/samples) by Novogene, Sacramento, CA. Stranded-mRNA seq was performed on Illumina Platform PE150 (Illumina, San Diego, CA) by Novogene at the UC Davis Sequencing Center.

Data analysis was performed as previously described (Laumet et al. 2015). Briefly, expression data of 4 samples per group were analyzed in R using bioconductor packages. STAR was used for alignment of paired-end reads to the mm10 version of the mouse reference genome feature. Counts was used to assign mapped sequence reads to genomic features, and DESeq2 was used to identify differentially expressed genes (padj<0.05). Qualify check of raw and aligned reads was performed with FastQC and Qualimap.

### Spectral flow cytometry

#### Single cell isolation from brain and spinal cord

Mice were euthanized using Carbon Dioxide and transcardially perfused with 20 ml 1X PBS. For brain and spinal cord single cell RNA isolation, both tissues were collected and placed in ice-cold 1X HBSS (Cat # 14-185-052; Gibco). Spine with attached muscle tissue was isolated from the body after the removal of the skin. The spinal cord from cervical vertebrae to lumbar vertebrae was collected via flushing 5 ml of PBS into the distal spinal column. Tissues were minced with either blade scalpel or scissors and then digested with 10 ml of enzyme mix: RPMI-1640 (Cat# R7509; Sigma,), 0.2 mg/ml DNase I (Cat# DN25; Sigma,), and 0.3 mg/ml Collagenase D (Cat# 11088858001; Sigma,). They were incubated for an hour at 37°C. After the incubation, tissues were transferred to a 70 µM Cell strainer and then smashed with plunger end of syringe. Ice-cold 1X HBSS was added during the procedure to minimize cell death. To inactivate the remaining enzymes, 4 ml of 10% FBS in 1X HBSS and 100 µl 0.5 M EDTA were added. After centrifuging them at 400 g, 4°C for 5 min, the pellet was resuspended with 4 ml of 37% Percoll (Cat# P1644; Sigma,). 4 ml of 70% Percoll was carefully underlaid using Pasteur pipette and 4 ml of 30% Percoll and 2 ml of 1X HBSS were overlaid. Following centrifugation (300 g, 40 min, 37°C, brake: 2), myelin layer on the top was removed and interface between 70% and 37% was collected and washed with 1X PBS.

#### Single cell isolation from brain and spinal meninge

For brain dura mater and spinal meninge single cell isolation, the dorsal part of the skull was carefully lifted from the brain. Spine was emptied by flushing 5 ml of PBS into the distal spinal column and then horizontally cut to expose the inner part. Under the stereoscope, the dura mater from the skull and the dura and leptomeninges from inner spine were peeled off from the skull and spine. Tissues were placed in ice-cold 1X HBSS. Following the centrifugation at 400 g, 4°C for 5 min, tissues were digested in 5 ml of enzyme cocktail for 30 minutes: RPMI-1640, 0.3 mg/ml DNase I, and 1 mg/ml Collagenase D. Every 10 min, the tubes with the tissues and enzyme were thoroughly inverted. After the incubation, tissues were transferred to a 70 µM Cell strainer and then smashed with plunger end of syringe. Ice-cold 1X HBSS was added during the procedure to minimize cell death. Then, they were centrifugated at 300 g, 4°C for 10 min, with minimal brake. The pellet was resuspended with 1X PBS.

#### Flow Cytometry

For staining of cell surface markers, cells were first stained with 1 μl/ml of LIVE/DEAD Blue (Cat# L34962; Invitrogen)-diluted in PBS for 15 min on the ice. Then, Fc-blocking antibodies CD16/CD32 (Cat# 553142; BD) were added to single-cell suspension before the surface staining to prevent non-specific binding ahead. Without washing, cells were stained with the surface marker antibodies, which are dissolved in FACS buffer (PBS, 1% BSA) and brilliant staining buffer (Cat# 563794; BD). Used antibodies are listed below. Cells were incubated at 4°C for 1 hour in the dark. Cells were analyzed with 5 lasers (16UV-16V-14B-10YG-8R) Cytek® Aurora - Spectral Flow Cytometry (Cytek, Fremont, CA). Ultracomp eBeadsTM (Cat# 01-2222-41; Invitrogen) labeled with each antibody were used as the reference controls for unmixing, except for eGFP. EGFP reference control was prepared from the spleen of the IL-10-eGFP mouse. Splenocytes without additional staining were analyzed by Cytek Aurora. Using SpectroFlo software, only the purified signature of eGFP from the eGFP positive splenocytes was exported as FCS file format and later imported as an eGFP reference control. The data were analyzed with SpectroFlo® software, FlowJo (TreeStar), and R studio.

To perform high-dimensional analysis, all data were imported into FlowJo, and only Live and CD45-positive cells were exported as FCS files. These files, containing the data of live-immune cells, were imported to RStudio for further analysis. Using R/Bioconductor packages, FlowSOM, and CATALYST, immune cells were computationally clustered based on their markers. The resulting data were processed for dimensionality reduction and visual representation with UMAP. Based on heatmaps and UMAP created from the analysis, we manually merged and annotated clusters. Through these steps, immune cell populations in the skin were defined computationally without supervision.

**Table 2.** Flow cytometry Antibody table.

| Antibody | Manufacturer | Catalog# | Location |
| --- | --- | --- | --- |
| LIVE/DEAD™ Fixable Blue Dead Cell Stain Kit | Invitrogen™ | L34962 | Carlsbad, CA |
| anti-CD45, BUV563 | BD Bioscience | 612924 | Franklin Lakes, NJ |
| anti-F4/80, APC-R700 | BD Bioscience | 565787 | Franklin Lakes, NJ |
| anti-CD11b, APC-Fire750 | BioLegend | 101261 | San Diego, CA |
| anti-Ly6G, BV711 | BD Bioscience | 563979 | Franklin Lakes, NJ |
| anti-CD11c, PE-Cy7 | BioLegend | 117317 | San Diego, CA |
| anti-CD19, BV785 | BioLegend | 115543 | San Diego, CA |
| anti-NK1.1, BUV805 | BD Bioscience | 741926 | Franklin Lakes, NJ |
| anti-CD3, APC-Fire810 | BioLegend | 100267 | San Diego, CA |
| anti-Ly6C, BV421 | BioLegend | 128031 | San Diego, CA |
| anti-FcεR1, Super bright 600 | Invitrogen™ | 63-589-882 | Carlsbad, CA |
| anti-SiglecF, Alexa fluor 647 | BioLegend | 155519 | San Diego, CA |
| anti-CD11c, BV650 | BioLegend | 117339 | San Diego, CA |
| anti-CD44, BUV395 | BD Bioscience | 740215 | Franklin Lakes, NJ |
| anti-CD206, Alexa Fluor 647 | BioLegend | 141712 | San Diego, CA |
| anti-Lyve1, PE | Invitrogen™ | 12-0443-82 | San Diego, CA |

**Table 3.** HSV vectors table.

| Name | Promoter | Catalog# | cDNAs |
| --- | --- | --- | --- |
| HSV-GFP (Ctrl) | hCMV | RN11 | GFP |
| HSV-mCherry-IRES-Cre | hCMV | RN91 | mCherry-IRES-Cre |
| HSV-mOprd1-IRES-mCherry | hCMV | Custom virus | mOprd1-IRES-mCherry |

hCMV: human cytomegalovirus, IRES: Internal ribosome entry site to allow cap independent translation.

## Statistical Analysis

Data are shown as mean ± standard error of the mean and the number of animals or samples used in each analysis are given in figure legends. GraphPad Prism 9 was used to analyze data for statistical tests, which are displayed in figure legends. Repeated measures two-way ANOVAs, one-way ANOVA, unpaired t-test, non-parametric Mann-Whitney tests, and Bonferroni’s correction for multiple tests were used based on experimental design. Statistical significances are indicated as follow * = p < 0.05, ** = p < 0.01, and *** = p < 0.001.

## Author contributions

Behaviors: KEI, GL, KBC; qPCR: KEI, KBC, KM, CME; Flow cytometry experiments and analysis: JS, YA, MB; RNA-seq: PB, JVM, GL. Drug/virus injection: KEI, JKF, GL; Mouse colony management: JFK, GL; Staining: KEI, GM, GS. Conceptualization: CJH, RD, AK, GL. Manuscript writing: KEI and GL.

## Acknowledgement

The present study was supported by the Rita Allen Foundation (G.L.), American Pain Society (G.L.), MSU Neuroscience Discretionary Research fund (GL), the NIH R01NS121259 (G.L. and Y.A.), R01NS073939 (C.J.H., R.D. and A.K.), and Department of Defense W81XWH-15-1-0076 (GS). GS is a New York Stem Cell Foundation _ Robertson Investigator.

We would like to thank Rachel Neve (Massachusetts General Hospital) for the construction of the HSV viral vectors.

## References

1. Bang, S., H. S. Kim, Y. S. Choo, S. W. Park, J. B. Chung and S. Y. Song (2006). “Differences in immune cells engaged in cell-mediated immunity after chemotherapy for far advanced pancreatic cancer.” Pancreas 32(1): 29–36.

2. Banimostafavi, E. S., M. Fakhar, S. Abediankenari, R. Alizadeh-Navaei, K. Mehdipour, V. Omrani-Nava and H. Majidi (2021). “Determining Serum Levels of IL-10 and IL-17 in Patients with Low Back Pain Caused by Lumbar Disc Degeneration.” Infect Disord Drug Targets 21(5): e270421185135.

3. Bardoni, R., V. L. Tawfik, D. Wang, A. François, C. Solorzano, S. A. Shuster, P. Choudhury, C. Betelli, C. Cassidy, K. Smith, J. C. de Nooij, F. Mennicken, D. O’Donnell, B. L. Kieffer, C. J. Woodbury, A. I. Basbaum, A. B. MacDermott and G. Scherrer (2014). “Delta opioid receptors presynaptically regulate cutaneous mechanosensory neuron input to the spinal cord dorsal horn.” Neuron 81(6): 1312–1327.

4. Basu, P., L. Custodio-Patsey, P. Prasoon, B. N. Smith and B. K. Taylor (2021). “Sex Differences in Protein Kinase A Signaling of the Latent Postoperative Pain Sensitization That Is Masked by Kappa Opioid Receptors in the Spinal Cord.” J Neurosci 41(47): 9827–9843.

5. Benkhart, E. M., M. Siedlar, A. Wedel, T. Werner and H. W. Ziegler-Heitbrock (2000). “Role of Stat3 in lipopolysaccharide-induced IL-10 gene expression.” J Immunol 165(3): 1612–1617.

6. Brackley, A. D., R. Gomez, A. N. Akopian, M. A. Henry and N. A. Jeske (2016). “GRK2 Constitutively Governs Peripheral Delta Opioid Receptor Activity.” Cell Rep 16(10): 2686–2698.

7. Bromberg, J. F., M. H. Wrzeszczynska, G. Devgan, Y. Zhao, R. G. Pestell, C. Albanese and J. E. Darnell, Jr. (1999). “Stat3 as an oncogene.” Cell 98(3): 295–303.

8. Bussmann, A. J. C., C. R. Ferraz, A. V. A. Lima, J. G. S. Castro, P. D. Ritter, T. H. Zaninelli, T. Saraiva-Santos, W. A. Verri, Jr. and S. M. Borghi (2022). “Association between IL-10 systemic low level and highest pain score in patients during symptomatic SARS-CoV-2 infection.” Pain Pract 22(4): 453–462.

9. Cahill, C. M., S. V. Holdridge, S. S. Liu, L. Xue, C. Magnussen, E. Ong, P. Grenier, A. Sutherland and M. C. Olmstead (2022). “Delta opioid receptor activation modulates affective pain and modality-specific pain hypersensitivity associated with chronic neuropathic pain.” J Neurosci Res 100(1): 129–148.

10. Campillo, A., D. Cabañero, A. Romero, P. García-Nogales and M. M. Puig (2011). “Delayed postoperative latent pain sensitization revealed by the systemic administration of opioid antagonists in mice.” Eur J Pharmacol 657(1-3): 89–96.

11. Chaplan, S. R., F. W. Bach, J. W. Pogrel, J. M. Chung and T. L. Yaksh (1994). “Quantitative assessment of tactile allodynia in the rat paw.” J Neurosci Methods 53(1): 55–63.

12. Chen, W., W. Walwyn, H. S. Ennes, H. Kim, J. A. McRoberts and J. C. Marvizón (2014). “BDNF released during neuropathic pain potentiates NMDA receptors in primary afferent terminals.” Eur J Neurosci 39(9): 1439–1454.

13. Chiang, A. C. A., X. Huo, A. Kavelaars and C. J. Heijnen (2019). “Chemotherapy accelerates age-related development of tauopathy and results in loss of synaptic integrity and cognitive impairment.” Brain Behav Immun 79: 319–325.

14. Corder, G., S. Doolen, R. R. Donahue, M. K. Winter, B. L. Jutras, Y. He, X. Hu, J. S. Wieskopf, J. S. Mogil, D. R. Storm, Z. J. Wang, K. E. McCarson and B. K. Taylor (2013). “Constitutive μ-opioid receptor activity leads to long-term endogenous analgesia and dependence.” Science 341(6152): 1394–1399.

15. Ding, Y., L. Qin, D. Zamarin, S. V. Kotenko, S. Pestka, K. W. Moore and J. S. Bromberg (2001). “Differential IL-10R1 expression plays a critical role in IL-10-mediated immune regulation.” J Immunol 167(12): 6884–6892.

16. Donnelly, R. P., H. Dickensheets and D. S. Finbloom (1999). “The interleukin-10 signal transduction pathway and regulation of gene expression in mononuclear phagocytes.” J Interferon Cytokine Res 19(6): 563–573.

17. Durante, M., S. Squillace, F. Lauro, L. A. Giancotti, E. Coppi, F. Cherchi, L. Di Cesare Mannelli, C. Ghelardini, G. Kolar, C. Wahlman, A. Opejin, C. Xiao, M. L. Reitman, D. K. Tosh, D. Hawiger, K. A. Jacobson and D. Salvemini (2021). “Adenosine A3 agonists reverse neuropathic pain via T cell-mediated production of IL-10.” J Clin Invest 131(7).

18. Faupel-Badger, J. M., L. C. R. Kidd, D. Albanes, J. Virtamo, K. Woodson and J. A. Tangrea (2008). “Association of IL-10 polymorphisms with prostate cancer risk and grade of disease.” Cancer Causes & Control 19(2): 119–124.

19. François, A. and G. Scherrer (2018). “Delta Opioid Receptor Expression and Function in Primary Afferent Somatosensory Neurons.” Handb Exp Pharmacol 247: 87–114.

20. Gaveriaux-Ruff, C., C. Nozaki, X. Nadal, X. C. Hever, R. Weibel, A. Matifas, D. Reiss, D. Filliol, M. A. Nassar, J. N. Wood, R. Maldonado and B. L. Kieffer (2011). “Genetic ablation of delta opioid receptors in nociceptive sensory neurons increases chronic pain and abolishes opioid analgesia.” Pain 152(6): 1238–1248.

21. Gendron, L., C. M. Cahill, M. von Zastrow, P. W. Schiller and G. Pineyro (2016). “Molecular Pharmacology of δ-Opioid Receptors.” Pharmacol Rev 68(3): 631–700.

22. Glocker, E. O., D. Kotlarz, C. Klein, N. Shah and B. Grimbacher (2011). “IL-10 and IL-10 receptor defects in humans.” Ann N Y Acad Sci 1246: 102–107.

23. Gonçalves, W. A., R. C. M. Ferreira, B. M. Rezende, G. A. B. Mahecha, M. Gualdron, F. H. P. de Macedo, I. D. G. Duarte, A. C. Perez, F. S. Machado, J. S. Cruz and T. R. L. Romero (2021). “Endogenous opioid and cannabinoid systems modulate the muscle pain: A pharmacological study into the peripheral site.” Eur J Pharmacol 901: 174089.

24. Grace, P. M., L. C. Loram, J. P. Christianson, K. A. Strand, J. G. Flyer-Adams, K. R. Penzkover, J. R. Forsayeth, A.-M. van Dam, M. J. Mahoney, S. F. Maier, R. A. Chavez and L. R. Watkins (2017). “Behavioral assessment of neuropathic pain, fatigue, and anxiety in experimental autoimmune encephalomyelitis (EAE) and attenuation by interleukin-10 gene therapy.” Brain, Behavior, and Immunity 59: 49–54.

25. Guo, Z., J. Zhang, X. Liu, J. Unsinger, R. S. Hotchkiss and Y. Q. Cao (2022). “Low-dose interleukin-2 reverses chronic migraine-related sensitizations through peripheral interleukin-10 and transforming growth factor beta-1 signaling.” Neurobiol Pain 12: 100096.

26. Hafezi, W., E. U. Lorentzen, B. R. Eing, M. Müller, N. J. King, B. Klupp, T. C. Mettenleiter and J. E. Kühn (2012). “Entry of herpes simplex virus type 1 (HSV-1) into the distal axons of trigeminal neurons favors the onset of nonproductive, silent infection.” PLoS Pathog 8(5): e1002679.

27. Hancock, M. J., C. M. Maher, P. Petocz, C. W. Lin, D. Steffens, A. Luque-Suarez and J. S. Magnussen (2015). “Risk factors for a recurrence of low back pain.” Spine J 15(11): 2360–2368.

28. Ho, A. S., Y. Liu, T. A. Khan, D. H. Hsu, J. F. Bazan and K. W. Moore (1993). “A receptor for interleukin 10 is related to interferon receptors.” Proceedings of the National Academy of Sciences 90(23): 11267–11271.

29. Inyang, K., M. D. Burton, T. Szabo-Pardi, E. Wentworth, T. A. McDougal, E. D. Ramirez, G. Pradhan, G. Dussor and T. Price (2019). “Indirect AMPK activators prevent incision-induced hyperalgesia and block hyperalgesic priming while positive allosteric modulators only block priming in mice.” J Pharmacol Exp Ther.

30. Inyang, K. E., S. R. George and G. Laumet (2021). “The µ-δ opioid heteromer masks latent pain sensitization in neuropathic and inflammatory pain in male and female mice.” Brain Res 1756: 147298.

31. Kabli, N. and C. M. Cahill (2007). “Anti-allodynic effects of peripheral delta opioid receptors in neuropathic pain.” Pain 127(1-2): 84–93.

32. Kandasamy, R., T. M. Hillhouse, K. E. Livingston, K. E. Kochan, C. Meurice, S. O. Eans, M. H. Li, A. D. White, B. P. Roques, J. P. McLaughlin, S. L. Ingram, N. T. Burford, A. Alt and J. R. Traynor (2021). “Positive allosteric modulation of the mu-opioid receptor produces analgesia with reduced side effects.” Proc Natl Acad Sci U S A 118(16).

33. Kohno, K., R. Shirasaka, K. Yoshihara, S. Mikuriya, K. Tanaka, K. Takanami, K. Inoue, H. Sakamoto, Y. Ohkawa, T. Masuda and M. Tsuda (2022). “A spinal microglia population involved in remitting and relapsing neuropathic pain.” Science 376(6588): 86–90.

34. Kotenko, S. V., C. D. Krause, L. S. Izotova, B. P. Pollack, W. Wu and S. Pestka (1997). “Identification and functional characterization of a second chain of the interleukin-10 receptor complex.” Embo j 16(19): 5894–5903.

35. Krukowski, K., N. Eijkelkamp, G. Laumet, C. E. Hack, Y. Li, P. M. Dougherty, C. J. Heijnen and A. Kavelaars (2016). “CD8+ T Cells and Endogenous IL-10 Are Required for Resolution of Chemotherapy-Induced Neuropathic Pain.” J Neurosci 36(43): 11074–11083.

36. Kühn, R., J. Löhler, D. Rennick, K. Rajewsky and W. Müller (1993). “Interleukin-10-deficient mice develop chronic enterocolitis.” Cell 75(2): 263–274.

37. Laumet, G., A. Bavencoffe, J. D. Edralin, X. J. Huo, E. T. Walters, R. Dantzer, C. J. Heijnen and A. Kavelaars (2020). “Interleukin-10 resolves pain hypersensitivity induced by cisplatin by reversing sensory neuron hyperexcitability.” Pain.

38. Laumet, G., J. D. Edralin, R. Dantzer, C. J. Heijnen and A. Kavelaars (2019). “Cisplatin educates CD8+ T cells to prevent and resolve chemotherapy-induced peripheral neuropathy in mice.” Pain 160(6): 1459–1468.

39. Laumet, G., J. Garriga, S. R. Chen, Y. Zhang, D. P. Li, T. M. Smith, Y. Dong, J. Jelinek, M. Cesaroni, J. P. Issa and H. L. Pan (2015). “G9a is essential for epigenetic silencing of K(+) channel genes in acute-to-chronic pain transition.” Nat Neurosci 18(12): 1746–1755.

40. Laumet, G., J. Ma, A. J. Robison, S. Kumari, C. J. Heijnen and A. Kavelaars (2019). “T Cells as an Emerging Target for Chronic Pain Therapy.” Front Mol Neurosci 12: 216.

41. Longoni, R., C. Cadoni, A. Mulas, G. Di Chiara and L. Spina (1998). “Dopamine-dependent behavioural stimulation by non-peptide delta opioids BW373U86 and SNC 80: 2. Place-preference and brain microdialysis studies in rats.” Behav Pharmacol 9(1): 9–14.

42. Madan, R., F. Demircik, S. Surianarayanan, J. L. Allen, S. Divanovic, A. Trompette, N. Yogev, Y. Gu, M. Khodoun, D. Hildeman, N. Boespflug, M. B. Fogolin, L. Gröbe, M. Greweling, F. D. Finkelman, R. Cardin, M. Mohrs, W. Müller, A. Waisman, A. Roers and C. L. Karp (2009). “Nonredundant roles for B cell-derived IL-10 in immune counter-regulation.” J Immunol 183(4): 2312–2320.

43. Maganin, A. G., G. R. Souza, M. D. Fonseca, A. H. Lopes, R. M. Guimarães, A. Dagostin, N. T. Cecilio, A. S. Mendes, W. A. Gonçalves, C. E. Silva, F. I. Fernandes Gomes, L. M. Mauriz Marques, R. L. Silva, L. M. Arruda, D. A. Santana, H. Lemos, L. Huang, M. Davoli-Ferreira, D. Santana-Coelho, M. B. Sant’Anna, R. Kusuda, J. Talbot, G. Pacholczyk, G. A. Buqui, N. P. Lopes, J. C. Alves-Filho, R. M. Leão, J. C. O’Connor, F. Q. Cunha, A. Mellor and T. M. Cunha (2022). “Meningeal dendritic cells drive neuropathic pain through elevation of the kynurenine metabolic pathway in mice.” J Clin Invest 132(23).

44. Marvizon, J. C., W. Walwyn, A. Minasyan, W. Chen and B. K. Taylor (2015). “Latent sensitization: a model for stress-sensitive chronic pain.” Curr Protoc Neurosci 71: 9.50.51-59.50.14.

45. McKelvey, R., T. Berta, E. Old, R. R. Ji and M. Fitzgerald (2015). “Neuropathic pain is constitutively suppressed in early life by anti-inflammatory neuroimmune regulation.” J Neurosci 35(2): 457–466.

46. Melemedjian, O. K., D. V. Tillu, J. K. Moy, M. N. Asiedu, E. K. Mandell, S. Ghosh, G. Dussor and T. J. Price (2014). “Local translation and retrograde axonal transport of CREB regulates IL-6-induced nociceptive plasticity.” Mol Pain 10: 45.

47. Milligan, E. D., K. R. Penzkover, R. G. Soderquist and M. J. Mahoney (2012). “Spinal Interleukin-10 Therapy to Treat Peripheral Neuropathic Pain.” Neuromodulation: Technology at the Neural Interface 15(6): 520–526.

48. Milligan, E. D., R. G. Soderquist, S. M. Malone, J. H. Mahoney, T. S. Hughes, S. J. Langer, E. M. Sloane, S. F. Maier, L. A. Leinwand, L. R. Watkins and M. J. Mahoney (2006). “Intrathecal polymer-based interleukin-10 gene delivery for neuropathic pain.” Neuron Glia Biol 2(4): 293–308.

49. Moore, K. W., R. de Waal Malefyt, R. L. Coffman and A. O’Garra (2001). “Interleukin-10 and the interleukin-10 receptor.” Annu Rev Immunol 19: 683–765.

50. Niehaus, J. K., B. Taylor-Blake, L. Loo, J. M. Simon and M. J. Zylka (2021). “Spinal macrophages resolve nociceptive hypersensitivity after peripheral injury.” Neuron 109(8): 1274–1282.e1276.

51. North, R. Y., Y. Li, P. Ray, L. D. Rhines, C. E. Tatsui, G. Rao, C. A. Johansson, H. Zhang, Y. H. Kim, B. Zhang, G. Dussor, T. H. Kim, T. J. Price and P. M. Dougherty (2019). “Electrophysiological and transcriptomic correlates of neuropathic pain in human dorsal root ganglion neurons.” Brain 142(5): 1215–1226.

52. Obara, I., W. Makuch, M. Spetea, J. Schütz, H. Schmidhammer, R. Przewlocki and B. Przewlocka (2007). “Local peripheral antinociceptive effects of 14-O-methyloxymorphone derivatives in inflammatory and neuropathic pain in the rat.” Eur J Pharmacol 558(1-3): 60–67.

53. Parisien, M., L. V. Lima, C. Dagostino, N. El-Hachem, G. L. Drury, A. V. Grant, J. Huising, V. Verma, C. B. Meloto, J. R. Silva, G. G. S. Dutra, T. Markova, H. Dang, P. A. Tessier, G. D. Slade, A. G. Nackley, N. Ghasemlou, J. S. Mogil, M. Allegri and L. Diatchenko (2022). “Acute inflammatory response via neutrophil activation protects against the development of chronic pain.” Sci Transl Med 14(644): eabj9954.

54. Pradhan, A., M. Smith, B. McGuire, C. Evans and W. Walwyn (2013). “Chronic inflammatory injury results in increased coupling of delta opioid receptors to voltage-gated Ca2+ channels.” Mol Pain 9: 8.

55. Prado, J., R. H. S. Westerink, J. Popov-Celeketic, C. Steen-Louws, A. Pandit, S. Versteeg, W. v. d. Worp, D. H. A. J. Kanters, K. A. Reedquist, L. Koenderman, C. E. Hack and N. Eijkelkamp (2021). “Cytokine receptor clustering in sensory neurons with an engineered cytokine fusion protein triggers unique pain resolution pathways.” Proceedings of the National Academy of Sciences 118(11): e2009647118.

56. Ray, P. R., S. Shiers, D. Tavares-Ferreira, I. Sankaranarayanan, M. L. Uhelski, Y. Li, R. Y. North, C. Tatsui, G. Dussor, M. D. Burton, P. M. Dougherty and T. J. Price (2021). “RNA Profiling of Neuropathic Pain-Associated Human DRGs Reveal Sex-differences in Neuro-immune Interactions Promoting Pain.” bioRxiv: 2021.2011.2027.470190.

57. Sabat, R., G. Grütz, K. Warszawska, S. Kirsch, E. Witte, K. Wolk and J. Geginat (2010). “Biology of interleukin-10.” Cytokine Growth Factor Rev 21(5): 331–344.

58. Scherrer, G., N. Imamachi, Y. Q. Cao, C. Contet, F. Mennicken, D. O’Donnell, B. L. Kieffer and A. I. Basbaum (2009). “Dissociation of the opioid receptor mechanisms that control mechanical and heat pain.” Cell 137(6): 1148–1159.

59. Scherrer, G., P. Tryoen-Tóth, D. Filliol, A. Matifas, D. Laustriat, Y. Q. Cao, A. I. Basbaum, A. Dierich, J. L. Vonesh, C. Gavériaux-Ruff and B. L. Kieffer (2006). “Knockin mice expressing fluorescent delta-opioid receptors uncover G protein-coupled receptor dynamics in vivo.” Proc Natl Acad Sci U S A 103(25): 9691–9696.

60. Shepherd, A. J., A. D. Mickle, J. P. Golden, M. R. Mack, C. M. Halabi, A. D. de Kloet, V. K. Samineni, B. S. Kim, E. G. Krause, R. W. Gereau and D. P. Mohapatra (2018). “Macrophage angiotensin II type 2 receptor triggers neuropathic pain.” Proceedings of the National Academy of Sciences 115(34): E8057–E8066.

61. Singh, S. K., K. Krukowski, G. O. Laumet, D. Weis, J. F. Alexander, C. J. Heijnen and A. Kavelaars (2022). “CD8+ T cell-derived IL-13 increases macrophage IL-10 to resolve neuropathic pain.” JCI Insight 7(5).

62. Sloane, E., S. Langer, B. Jekich, J. Mahoney, T. Hughes, M. Frank, W. Seibert, G. Huberty, B. Coats, J. Harrison, D. Klinman, S. Poole, S. Maier, K. Johnson, R. Chavez, L. R. Watkins, L. Leinwand and E. Milligan (2009). “Immunological priming potentiates non-viral anti-inflammatory gene therapy treatment of neuropathic pain.” Gene Therapy 16(10): 1210–1222.

63. Song, X. J., Z. J. Huang, W. B. Song, X. S. Song, A. F. Fuhr, A. L. Rosner, H. Ndtan and R. L. Rupert (2016). “Attenuation Effect of Spinal Manipulation on Neuropathic and Postoperative Pain Through Activating Endogenous Anti-Inflammatory Cytokine Interleukin 10 in Rat Spinal Cord.” J Manipulative Physiol Ther 39(1): 42–53.

64. Spina, L., R. Longoni, A. Mulas, K. J. Chang and G. Di Chiara (1998). “Dopamine-dependent behavioural stimulation by non-peptide delta opioids BW373U86 and SNC 80: 1. Locomotion, rearing and stereotypies in intact rats.” Behav Pharmacol 9(1): 1–8.

65. Szabo-Pardi, T. A., L. R. Barron, M. E. Lenert and M. D. Burton (2021). “Sensory Neuron TLR4 mediates the development of nerve-injury induced mechanical hypersensitivity in female mice.” Brain Behav Immun 97: 42–60.

66. Takeda, K., B. E. Clausen, T. Kaisho, T. Tsujimura, N. Terada, I. Förster and S. Akira (1999). “Enhanced Th1 activity and development of chronic enterocolitis in mice devoid of Stat3 in macrophages and neutrophils.” Immunity 10(1): 39–49.

67. Taylor, B. K. and G. Corder (2014). “Endogenous analgesia, dependence, and latent pain sensitization.” Curr Top Behav Neurosci 20: 283–325.

68. Taylor, B. K., G. P. Sinha, R. R. Donahue, C. M. Grachen, J. A. Morón and S. Doolen (2019). “Opioid receptors inhibit the spinal AMPA receptor Ca(2+) permeability that mediates latent pain sensitization.” Exp Neurol 314: 58–66.

69. Uçeyler, N., J. P. Rogausch, K. V. Toyka and C. Sommer (2007). “Differential expression of cytokines in painful and painless neuropathies.” Neurology 69(1): 42–49.

70. Vanderwall, A. G., S. Noor, M. S. Sun, J. E. Sanchez, X. O. Yang, L. L. Jantzie, N. Mellios and E. D. Milligan (2018). “Effects of spinal non-viral interleukin-10 gene therapy formulated with d-mannose in neuropathic interleukin-10 deficient mice: Behavioral characterization, mRNA and protein analysis in pain relevant tissues.” Brain, Behavior, and Immunity 69: 91–112.

71. Wang, D., V. L. Tawfik, G. Corder, S. A. Low, A. François, A. I. Basbaum and G. Scherrer (2018). “Functional Divergence of Delta and Mu Opioid Receptor Organization in CNS Pain Circuits.” Neuron 98(1): 90–108.e105.

72. Wang, H., C. J. Heijnen, C. T. van Velthoven, H. L. Willemen, Y. Ishikawa, X. Zhang, A. K. Sood, A. Vroon, N. Eijkelkamp and A. Kavelaars (2013). “Balancing GRK2 and EPAC1 levels prevents and relieves chronic pain.” J Clin Invest 123(12): 5023–5034.

73. Wang, W. W., Y. Wang, K. Li, R. Tadagavadi, W. E. Friedrichs, M. Budatha and W. B. Reeves (2020). “IL-10 from dendritic cells but not from T regulatory cells protects against cisplatin-induced nephrotoxicity.” PLoS One 15(9): e0238816.

74. Wangzhou, A., C. Paige, S. V. Neerukonda, D. K. Naik, M. Kume, E. T. David, G. Dussor, P. R. Ray and T. J. Price (2021). “A ligand-receptor interactome platform for discovery of pain mechanisms and therapeutic targets.” Sci Signal 14(674).

75. Watkins, L. R., R. A. Chavez, R. Landry, M. Fry, S. M. Green-Fulgham, J. D. Coulson, S. D. Collins, D. K. Glover, J. Rieger and J. R. Forsayeth (2020). “Targeted interleukin-10 plasmid DNA therapy in the treatment of osteoarthritis: Toxicology and pain efficacy assessments.” Brain Behav Immun 90: 155–166.

76. Weber-Nordt, R. M., J. K. Riley, A. C. Greenlund, K. W. Moore, J. E. Darnell and R. D. Schreiber (1996). “Stat3 Recruitment by Two Distinct Ligand-induced, Tyrosine-phosphorylated Docking Sites in the Interleukin-10 Receptor Intracellular Domain*.” Journal of Biological Chemistry 271(44): 27954–27961.

77. Wei, Z., Y. Fei, W. Su and G. Chen (2019). “Emerging Role of Schwann Cells in Neuropathic Pain: Receptors, Glial Mediators and Myelination.” Front Cell Neurosci 13: 116.

78. Willemen, H., A. Kavelaars, J. Prado, M. Maas, S. Versteeg, L. J. J. Nellissen, J. Tromp, R. Gonzalez Cano, W. Zhou, M. E. Jakobsson, J. Małecki, G. Posthuma, A. M. Habib, C. J. Heijnen, P. Falnes and N. Eijkelkamp (2018). “Identification of FAM173B as a protein methyltransferase promoting chronic pain.” PLoS Biol 16(2): e2003452.

79. Zeisel, A., H. Hochgerner, P. Lönnerberg, A. Johnsson, F. Memic, J. van der Zwan, M. Häring, E. Braun, L. E. Borm, G. La Manno, S. Codeluppi, A. Furlan, K. Lee, N. Skene, K. D. Harris, J. Hjerling-Leffler, E. Arenas, P. Ernfors, U. Marklund and S. Linnarsson (2018). “Molecular Architecture of the Mouse Nervous System.” Cell 174(4): 999–1014.e1022.

80. Zelaya, C. E., J. M. Dahlhamer, J. W. Lucas and E. M. Connor (2020). “Chronic Pain and High-impact Chronic Pain Among U.S. Adults, 2019.” NCHS Data Brief(390): 1-8.

81. Zorina-Lichtenwalter, K., M. Parisien and L. Diatchenko (2018). “Genetic studies of human neuropathic pain conditions: a review.” Pain 159(3): 583–594.

