## Supplemental methods for "Tonic Meningeal Interleukin-10 Upregulates Delta Opioid Receptor to Prevent Relapse to Pain"

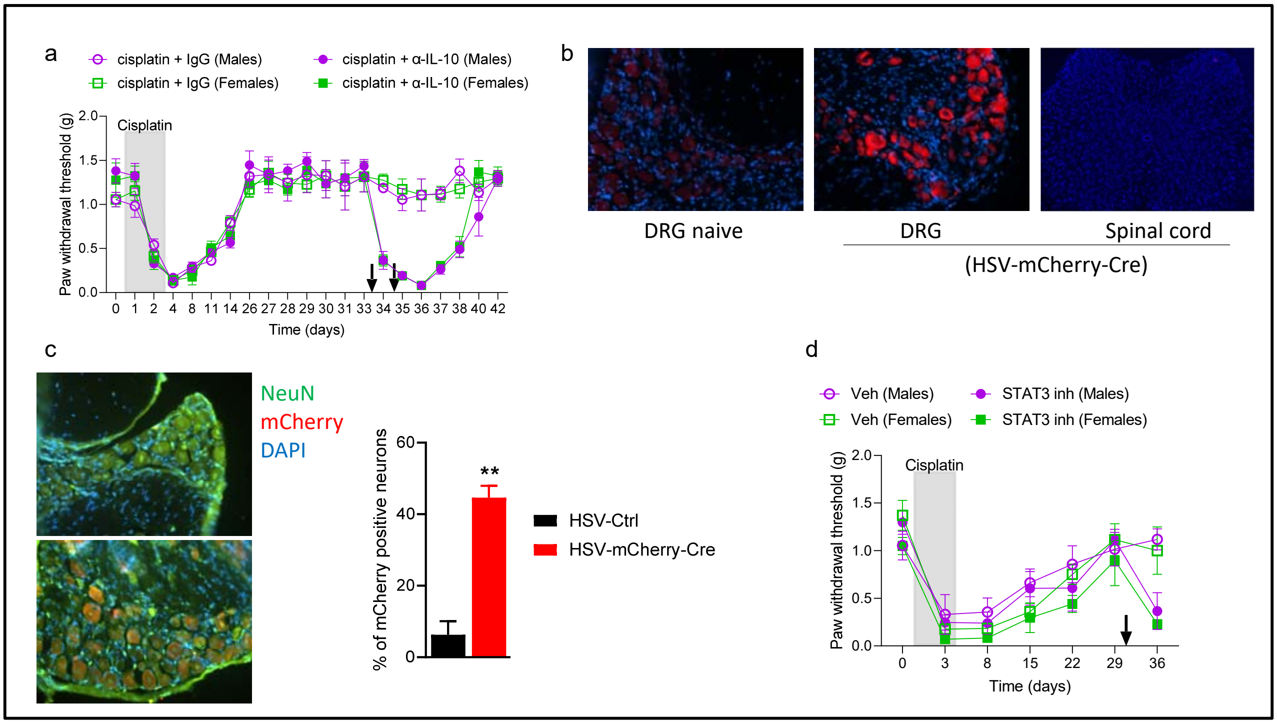


**Supplementary figure 1. Inhibition of IL-10-STAT3 signaling reinstates pain hypersensitivity in both sexes**. **a**, cisplatin administration (2mg/kg/day x3, gray area) induces mechanical hypersensitivity measured by von Frey filaments. Intrathecal injection of neutralizing anti-IL-10 antibody reinstates mechanical sensitivity in both male and female mice (n=5/group). **b**, Representative images of mCherry staining in DRG and spinal cord sections in naïve untreated mice and HSV-mCherry-Cre treated mice. **c**, Representative images of DRG sections of naïve and HSV-mCherry-Cre vector-treated mice. DRG sections are stained with anti-NeuN (green), mCherry (vector), and DAPI. Quantification of the percentages of double positive (NeuN+ mCherry+) cells. Unpaired t-test, $p<0.002$ (n=3/group). **d**, STAT3 inhibitor (inh) BP-1-102 (1 mg/kg) reinstates mechanical hypersensitivity in both cisplatin-treated male and female mice (n=5/group).

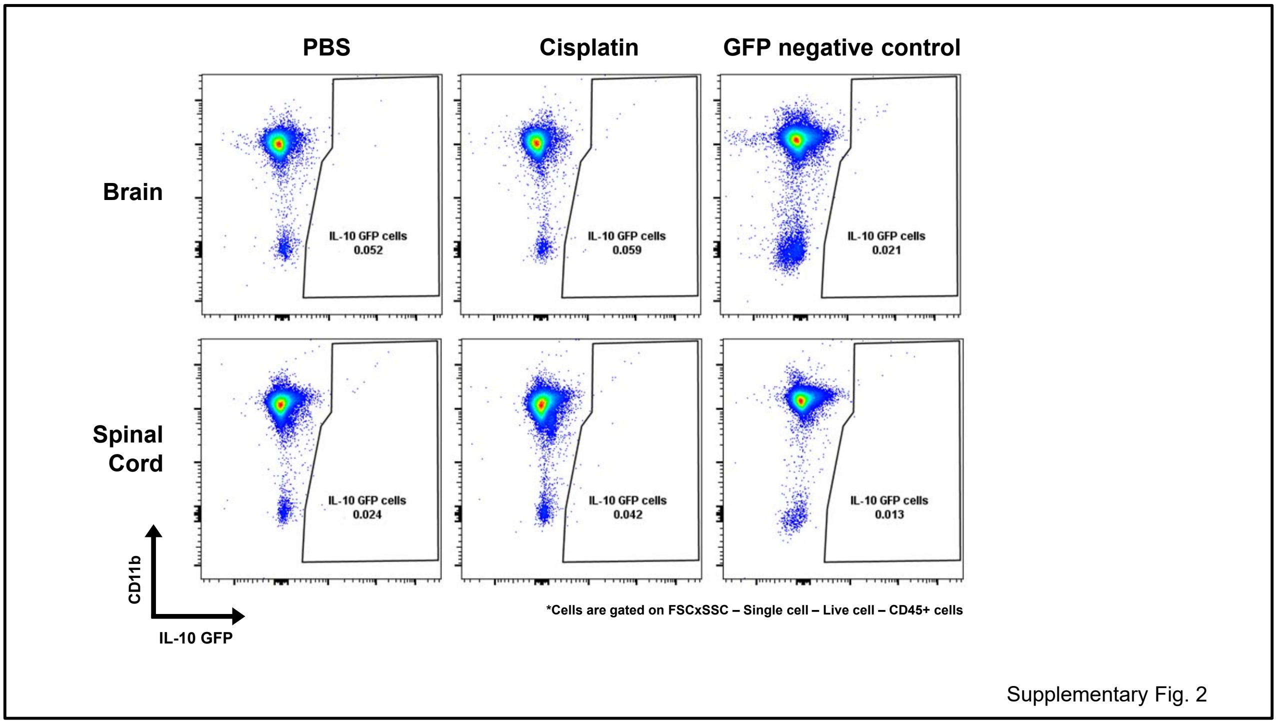


**Supplementary figure 2. No GFP+ (IL-10 producing) cells were detected in the brain and spinal cord parenchyma by flow cytometry**. Flow cytometry was performed from PBS- and cisplatin-treated IL-10 GFP reporter mice from both sexes. C57Bl6 WT mice were used as negative control (no expression of GFP) to set up the GFP gating.

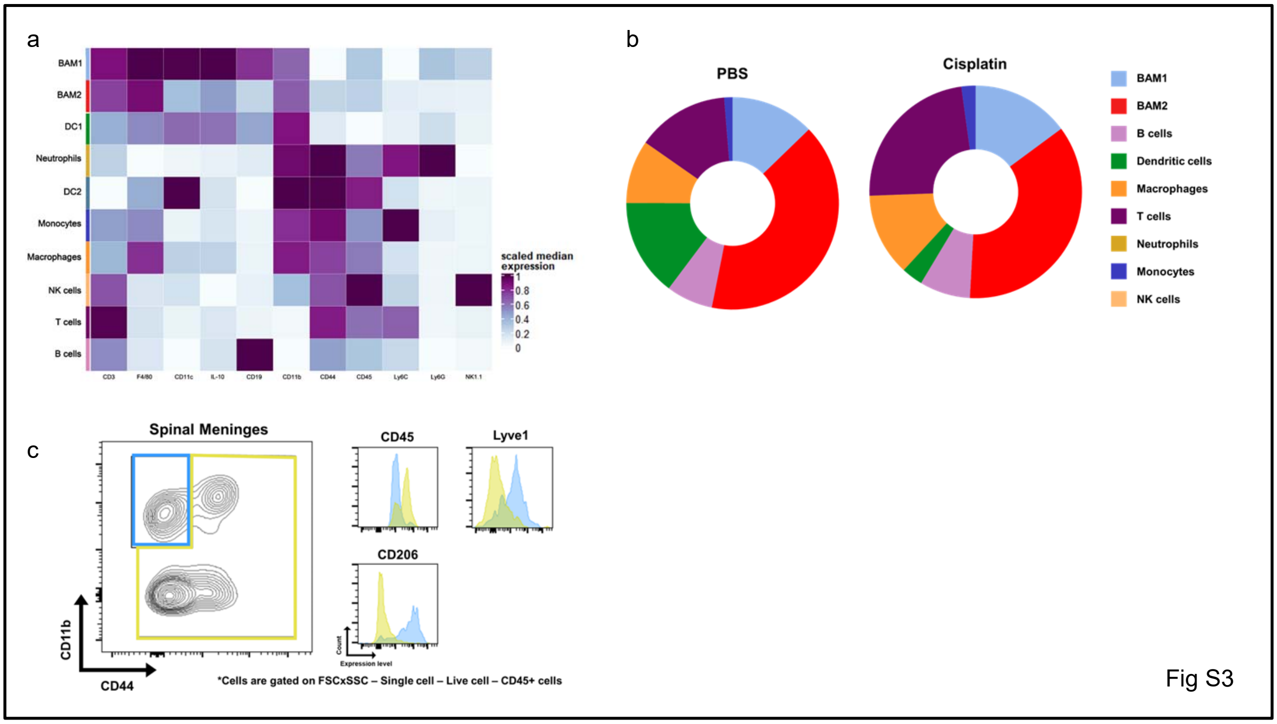


**Supplementary figure 3. IL-10 is produced by meningeal resident macrophages**. **a**, heat map shows the unsupervised clustering of immune cells based on different markers. Meningeal resident macrophages or border-associated macrophages (BAMs) are identified as CD45low, CD11b+, F4/80+, CD44-, Ly6C-. **b**, Pie chart of IL-10 producing meningeal immune cell population. **c**, CD11b+, CD44(-) cells are CD45low, Lyve1+, CD206+ confirming that resident macrophages/BAMs identity.

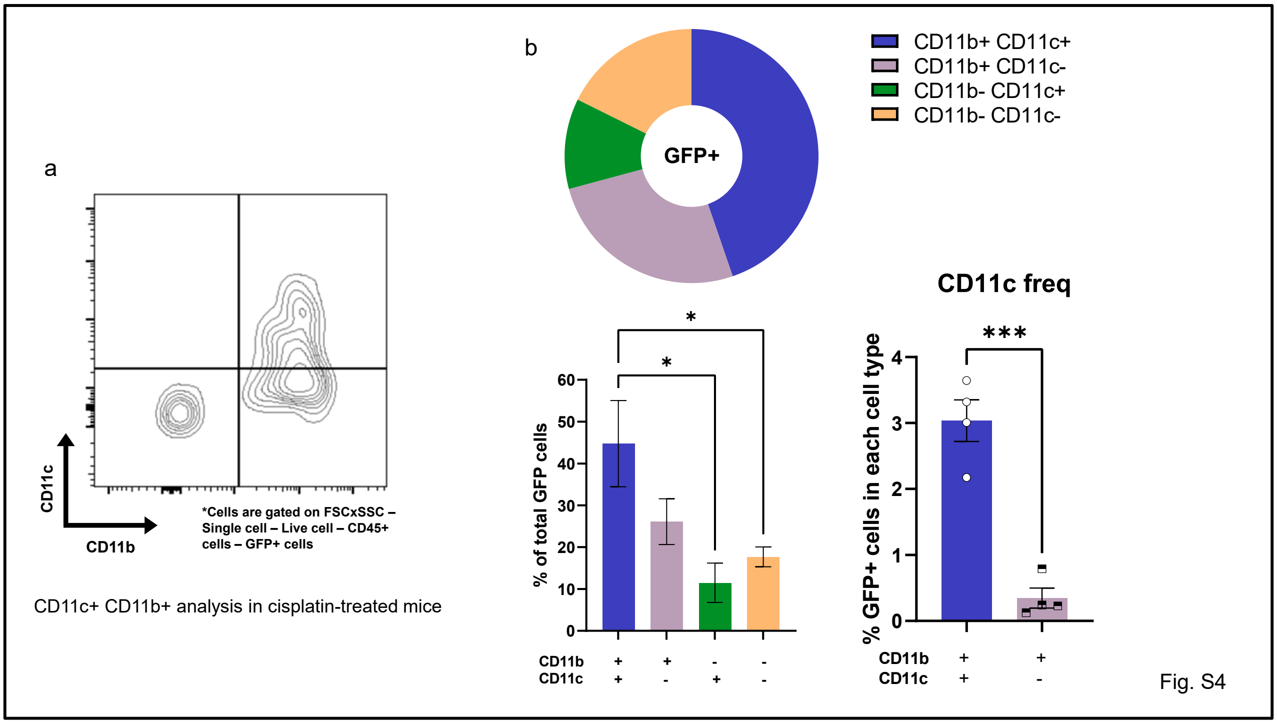


**Supplementary figure 4. IL-10 is produced by CD11c+ macrophages**. **a**, Representative flow plot of CD45+GFP+ cells assessed for CD11b and CD11c markers. **b**, CD11b+CD11c+ cells are more important producers of IL-10 than CD11b+CD11c(-) cells. (n=4/group)

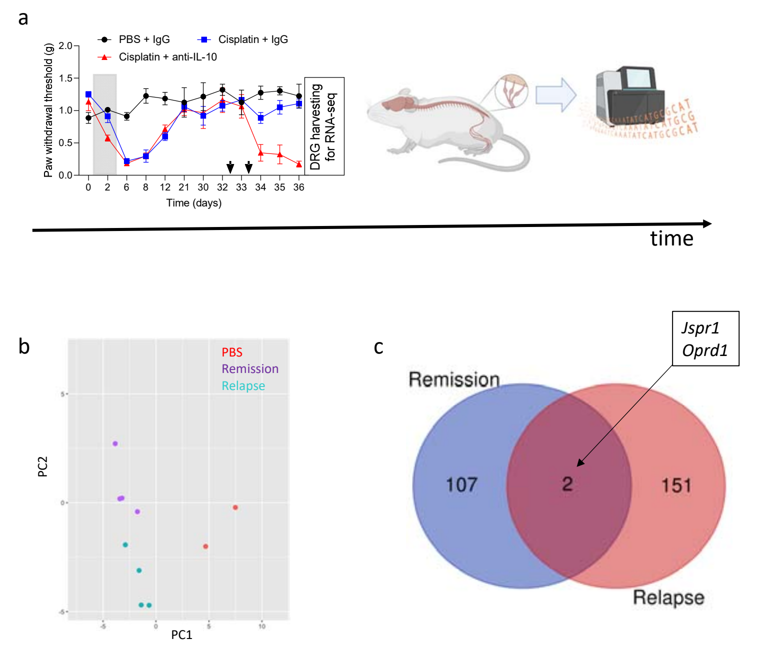


Suppl Figure S5

**Supplementary figure 5. Remission and Relapse transcriptomic signature . a**, Experimental design for DRG RNA-sequencing to assess the transcriptome associated with Remission and Relapse of neuropathic pain. **b**, Principal Component Analysis (PCA) shows that DRG transcriptome from mice in Remission and Relapse cluster separately. **c**, Venn diagram shows that only 2 common genes differentially expressed in Remission and Relapse; *Jspr1* and *Oprd1*.

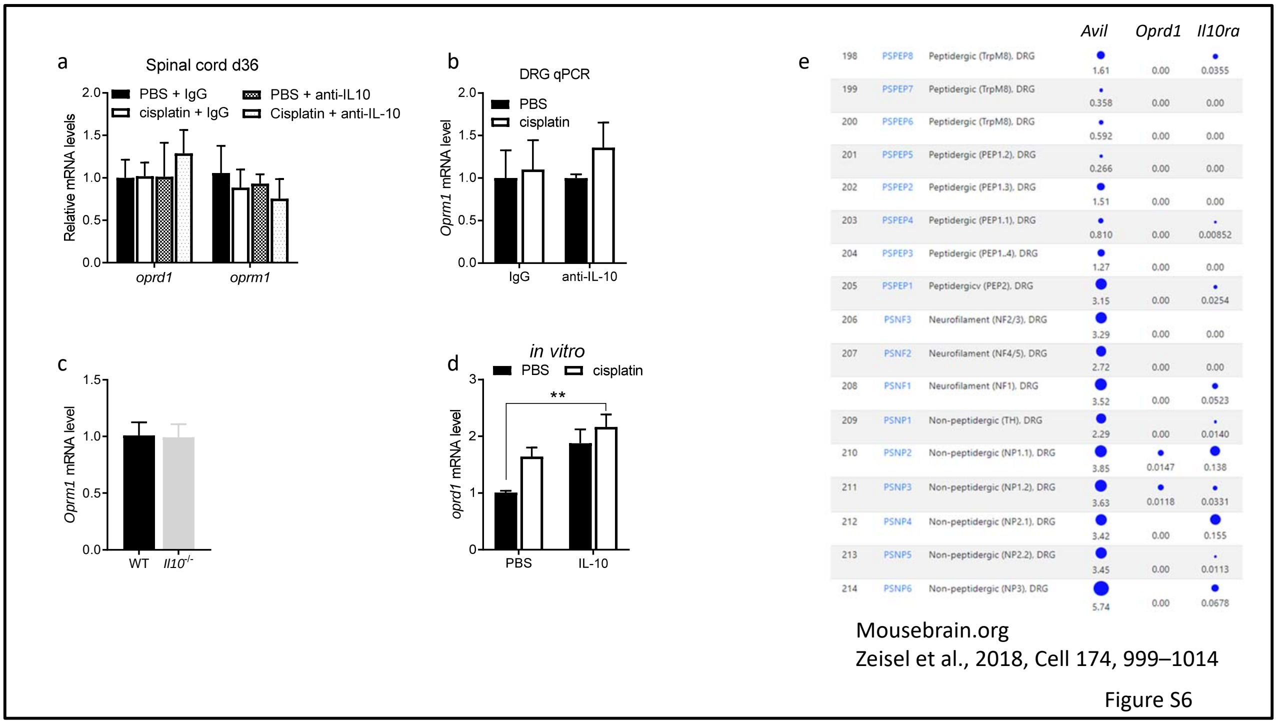


**Supplementary figure 6. IL-10 signaling does not regulate the expression of *Oprm1* in the DRG neither *Oprd1* in the spinal cord**. **a**, *Oprd1* and *Oprm1* expression in the spinal cord are not affected by cisplatin or IL-10 signaling inhibition on day 36 after PBS or cisplatin administration (n=5-7/group). **b**, *Oprm1* mRNA levels in the DRG is not affected by anti-IL-10 on day 36 after cisplatin (Remission) (n=4-6/group). **c**, *Oprm1* mRNA levels in the DRG is not affected by the lack of IL-10 after cisplatin (n=10-11/group). **d**, *Oprd1* mRNA levels is increased in DRG cell line 50B11 cells in response to cisplatin and recombinant IL-10 (10 ng/ml, Biovision cat#4156-50) (n=7/group). Two-way ANOVA IL-10 effect F(1,24) = 14.3, p=0.0009. **e**, Snapshot from mousebrain.org (Zeisel et al., 2018) shows that *Oprd1* and *Il10ra* are expressed by similar cells in the DRG.

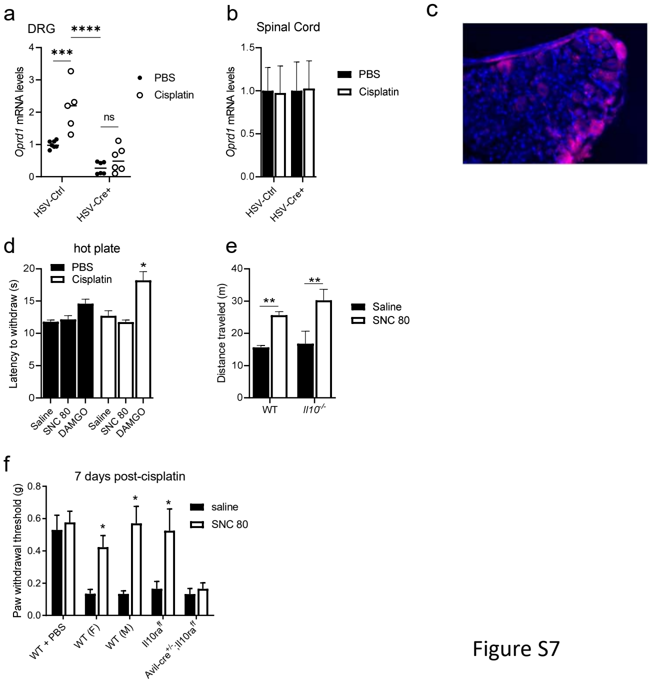


Figure S7

**Supplementary figure 7. Behavioral effects of δOR during remission**. **a**, *Oprd1* expression in DRG is drastically decreased in *Oprd1*$^{floxflox}$ mice treated with HSV-mCherry-Cre (n=5-7/group), Two-way ANOVA Cre effect F(1,20) = 49.3, p<0.0001. **b**, Intraplantar injection of HVS-mCherry-Cre vector does not affect *Oprd1* expression in *Oprd1*$^{floxflox}$ mice. **c**, Representative image of DRG section with mCherry positive cells. **d**, Analgesic effect of MOR agonist DAMGO on the hot plate test (55$^{o}$C) is more pronounced in mice in remission (day 36 after cisplatin). δOR agonist SNC 80 (10 mg/kg) has no effect of the hot plate (n=6/group), Two-way ANOVA followed by Tukey's correction for multiple tests: DAMGO-PBS vs. DAMGO-cisplatin p=0.022. **e**, SNC 80-induced increased locomotor activity is not affected by IL-10 signaling (n=4-7/group), Two-way ANOVA drug effect F(1,17) = 30.5, p<0.0001. **f**, SNC 80 reversed mechanical hypersensitivity 7 days after cisplatin in both sexes, but SNC 80 does not affect mechanical hypersensitivity in mice that lack IL-10R1 on DRG advillin-positive neurons (n=4-6/group), * indicates p<0.05 for comparison saline vs SNC80 in the same group analyzed by twoway ANOVA followed by Bonferroni's correction for multiple tests.

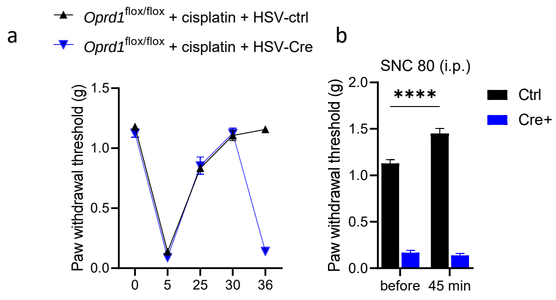


Figure S8

**Supplementary figure 8. DRG *Oprd1* expression is necessary for SNC80-induced analgesia**. **a**, Intraplantar injection of HSV-mCherry-Cre (day 32) in cisplatin-treated *Oprd1*$^{floxflox}$ mice reinstated mechanical hypersensitivity (n=6-7/group). **b**, injection of SNC 80 (10 mg/kg) does not increase mechanical sensitivity in mice lacking *Oprd1* in DRG neurons. Two-way ANOVA Cre x SNC80 interaction $F(1,22) = 26.6$, $p<0.0001$.
